# Energy Landscape Analysis with Automated Region-of-Interest Selection via Genetic Algorithms

**DOI:** 10.1101/2025.08.05.668814

**Authors:** Koichiro Mori, Tomoyuki Hiroyasu, Satoru Hiwa

## Abstract

Understanding brain dynamics is essential for advancing cognitive and clinical neuroscience. Energy landscape analysis (ELA), based on the pairwise maximum entropy model, is a powerful framework to characterize brain activity as transitions among discrete states defined by regional activity patterns. However, traditional ELA relies on the subjective manual selection of a small subset of regions of interest (ROIs) from whole-brain parcellations to satisfy mathematical constraints, which limits the scope and reproducibility of ELA results due to subjective subset selection. To overcome this, we developed ELA/GAopt, a meta-framework that utilizes a genetic algorithm to automate the selection of ROI combinations from the entire atlas search space by optimizing a user-defined objective function. In this study, we implemented a representative objective function balancing model fitting accuracy with the inter-individual variability of model parameters. We applied ELA/GAopt to three independent resting-state functional magnetic resonance imaging datasets. In Scenario 1, using the Creativity dataset (OpenNeuro: ds002330, *n* = 61), the ROI sets identified by ELA/GAopt achieved significantly higher objective function values and pattern reproducibility than randomly selected ROI sets (*p* < 0.05). Additional validation with the large-scale Human Connectome Project Young Adult (HCP-YA) dataset (*n* = 270) confirmed the robustness of our framework in high-dimensional settings. Stability analysis using Jaccard and Hamming metrics demonstrated that ELA/GAopt consistently identified reproducible ROI subsets across independent optimization runs. In Scenarios 2-4, we analyzed data from the Autism Brain Imaging Data Exchange II dataset using a site-disjoint validation design to mitigate findings were robust against multi-site artifacts. ELA/GAopt identified ASD-specific dynamics, where participants tend to visit local minima characterized by global co-activation of selected ROIs within sensory-motor and visual networks. These signatures were replicated in an independent cohort consisting of different scanning sites. Furthermore, ROI sets optimized for one group (ASD or typically developing controls) did not generalize to the other, highlighting distinct neurodynamic architectures. These results demonstrate that ELA/GAopt provides a reproducible, data-driven pathway for characterizing condition-specific brain dynamics, serving as a methodological basis for future, harmonization-aware and externally validated biomarker studies.

## Introduction

Interpreting system behavior from multidimensional observation data is essential in various scientific fields, including neuroscience. Energy landscape analysis (ELA) is one such method that applies the Ising model to multidimensional data, introducing the concept of energy to each pattern and visualizing the stability and transitions of emerging patterns as an energy landscape. The Ising model, also known as the pairwise maximum entropy model (pMEM) (Saberi et al., 2024; Watanabe et al., 2013), was initially introduced in statistical physics to explain the properties of magnetic materials. It is highly versatile and facilitates the interpretation of system dynamics as interactions between paired elements. The energy landscape derived from observed multidimensional data enables a bird’s-eye view of the system’s dynamics. A notable application in neuroscience was introduced by Watanabe, Hirose, et al. (2014), who applied the ELA to functional MRI (fMRI) data, establishing a data-driven approach to extract brain dynamics from neuroimaging data. Subsequently, Ezaki et al. (2017) investigated fundamental issues of ELA for neuroimaging data and developed the Energy Landscape Analysis Toolbox (ELAT). Their contributions have made ELA applicable to diverse neuroimaging datasets, including various neurological conditions and disorders (Saberi et al., 2024; Xing et al., 2024; Yamashita et al., 2021), and significantly advanced understanding in the neuroscience field (Ezaki et al., 2017, 2018; Hosaka et al., 2025; Jeong et al., 2021; Kang et al., 2019, 2021; Kondo et al., 2022; Saberi et al., 2024; Watanabe, Hirose, et al., 2014; Watanabe, Masuda, et al., 2014; Watanabe & Rees, 2017; Xing et al., 2024).

Despite its success, the current ELA paradigm faces several challenges due to its analytical constraints, including the length of data and the selection of the region of interest (ROI). The ELA requires sufficient data length to ensure the accurate fitting of the pMEM, which necessitates an exponential increase in the required data length with the number of variables, roughly doubling for each additional variable. Masuda et al. (2025) estimated that the practical upper limit for the number of ROIs (i.e., data dimension) that can be included in ELA without compromising accuracy is around 10 to 15, depending on the available data length (Masuda et al., 2025).

In contrast, whole-brain parcellation typically yields hundreds of regions, depending on the atlas used (e.g., 116 for the AAL (Tzourio-Mazoyer et al., 2002), 264 for the Power atlas (Power et al., 2011), and 100 to 1000 for the Schaefer 2018 atlas (Schaefer et al., 2018; Yeo et al., 2011)). This creates a significant gap between the available anatomical resolution (hundreds of regions provided by atlases) and the upper limit imposed by ELA’s mathematical constraints (approximately 10-15 regions mentioned above). Ideally, to find the system that accurately represents brain dynamics, all possible combinations of regions should be analyzed; however, selecting 10-15 ROIs from hundreds leads to a combinatorial explosion, making an exhaustive search impossible. Due to this constraint, traditional studies have relied on manual subset selection, such as selecting representative regions involved in relevant cognitive processes based on prior research or choosing representative regions from known functional networks. While these methods utilize standard atlases, the choice of the ROI subset often remains subjective and hypothesis-driven, which can hinder a more systematic, data-driven exploration of ELA outcomes.

To overcome these limitations, we propose ELA/GAopt, a genetic algorithm-based optimization framework designed to automate ROI selection in ELA while ensuring model fitting accuracy. Unlike traditional hypothesis-driven selection, our approach treats the entire brain atlas as a search space to identify an optimal combination of ROIs that satisfies the mathematical assumptions of pMEM. Genetic algorithms (GAs) are a class of heuristic optimization techniques inspired by natural selection and evolution. They are widely used to obtain approximate solutions for nonlinear optimization problems, and particularly suited for this task due to their ability to navigate complex, non-linear search spaces through population-based heuristic exploration, allowing for the discovery of interactions that simple greedy methods such as recursive feature elimination might overlook (Bulut et al., 2025; Vafaie & Imam, n.d.). In our proposed framework, ROI selection is formulated as a combinatorial optimization problem where the objective function can be flexibly defined according to specific research goals. As a representative example in this study, we employ an objective function that incorporates both pMEM fitting accuracy and the inter-individual variability of model parameters. This procedure eliminates the subjective bias inherent in manual ROI selection, enabling a more systematic implementation of ELA—a critical advancement for automated exploration of condition-specific brain dynamics.

The primary goal of this study is to establish ELA/GAopt as a methodological framework that resolves ELA’s dimensional constraints while preserving anatomical interpretability. Unlike the conventional methods such as independent component analysis (ICA)-based parcellations, clustering approaches, and machine learning-driven feature selection, our focus is on identifying ‘analysis-ready’ ROI subsets specifically optimized for downstream energy landscape modeling. To validate the effectiveness and robustness of our approach, we evaluate it using three open neuroimaging datasets. First, we assess its generalizability by testing whether the brain state dynamics identified using the proposed method are reproducible on the dataset of healthy participants using both the Creativity dataset and the Human Connectome Project Young Adult (HCP-YA) dataset. Then, we investigate its clinical applicability by analyzing resting-state fMRI data from the Autism Brain Imaging Data Exchange II (ABIDE II). This analysis enables us to investigate whether the method can identify condition-specific patterns in brain state transitions between individuals with autism spectrum disorder (ASD) and typically developing controls (CTL). These applications demonstrate the potential of ELA/GAopt to enhance the objectivity and reliability of brain dynamics modeling, thereby contributing to provide analysis-ready ROI subsets under pMEM constraints.

## Methods

### Overview of ELA/GAopt: Energy landscape analysis with automated ROI selection via genetic algorithms

The proposed ELA/GAopt framework aims to automate the selection of ROIs in ELA through a combinatorial optimization approach. It involves iteratively selecting ROI subsets, performing ELA, particularly fitting the pMEM, and evaluating the results until specified conditions are met.

To solve this combinatorial optimization problem, we employ a GA. GAs are heuristic optimization techniques characterized by multipoint exploration enabling the generation of multiple candidate solutions (i.e., ROI subsets) in parallel. Each solution is evaluated using a fitness function that quantifies the goodness of the solution, which is based on the ELA output in this case. This strategy helps avoid local optima and improves the robustness of the search process.

A schematic overview of ELA/GAopt in neuroimaging data is presented in Figure 1. The following sections describe each component of the framework and detailed analysis procedures.

**Figure 1.**
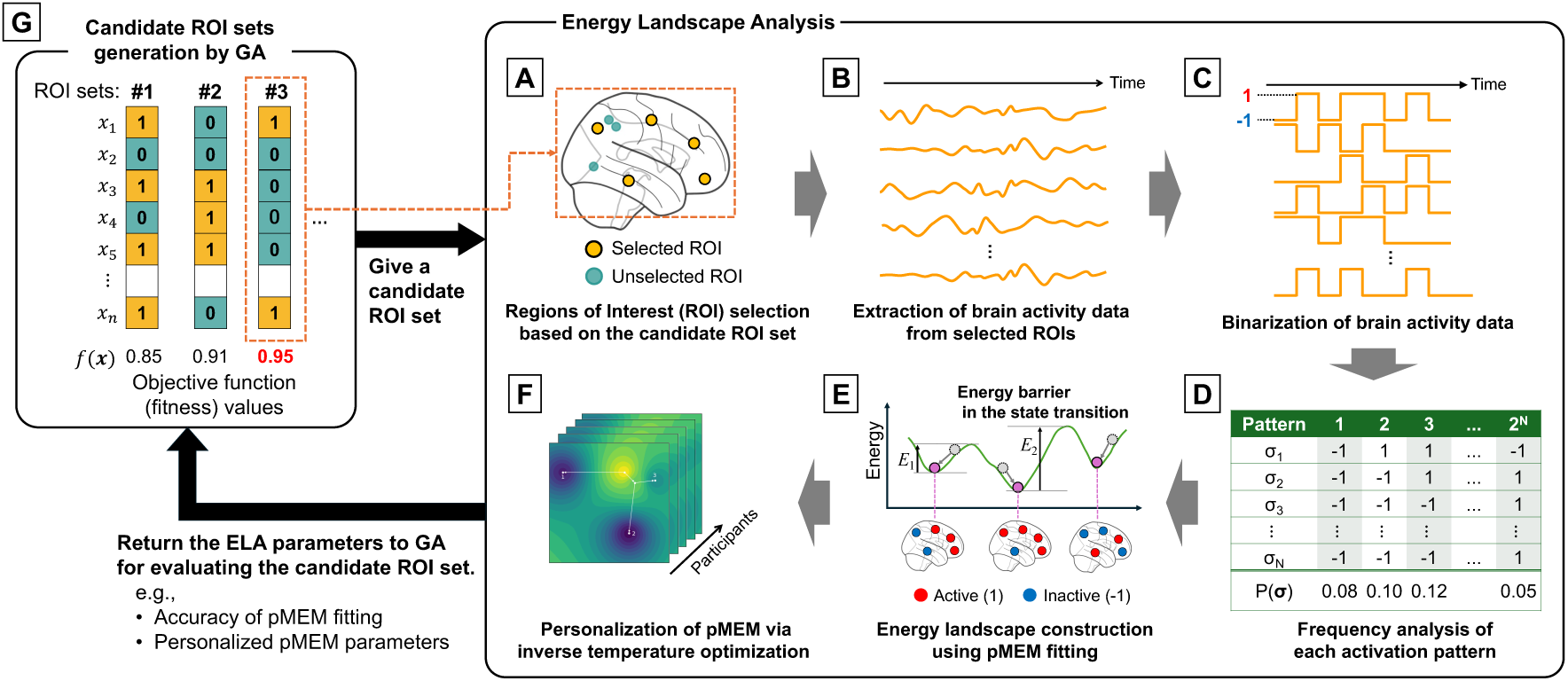
Schematic overview of ELA/GAopt framework. The proposed framework ELA/GAopt combines energy landscape analysis (ELA) with a genetic algorithm (GA)-based optimization for automatic region of interest (ROI) selection. (**A**) Candidate ROI sets are generated as binary vectors by the GA, where each gene indicates whether a specific ROI is selected (1) or not (0). (**B**) Brain activity data are extracted from the selected ROIs. (**C**) The brain activity data are binarized (active: +1 or inactive: −1) to form discrete activation patterns over time. (**D**) The frequency of occurrence of each available activation pattern is analyzed. (**E**) A pairwise maximum entropy model (pMEM) is fitted to the binarized time series data to estimate the energy associated with each observed signal pattern. The resulting energy landscape visualizes the probability and energy of each activation pattern, characterizing brain dynamics. (**F**) A two-step pMEM parameter optimization method, in which the inverse temperature parameter—which defines the Boltzmann distribution—is optimized for each individual dataset while keeping the group-derived pMEM parameters fixed. (**G**) The ELA-derived parameters (e.g., model accuracy and personalized parameters) are fed back into the GA as a fitness score to evaluate each candidate ROI set. The GA then iteratively updates the ROI sets through selection, crossover, mutation, and repair operations.

### Framework of ELA

#### Binarization of multidimensional time series data

The input to ELA is an *N*-dimensional discrete time series obtained from *N* ROIs:

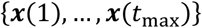

where ***x***(*t*) ∈ ℝ^*N*^ for *t* ∈ {1, …, *t*_max_} is the ROI signal vector at time *t*, and *t*_max_ is the total number of time points (i.e., the length of the observed time series), Each vector ***x***(*t*) is defined as:

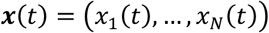

In ELA, multidimensional signals at each time point are converted into binary patterns. Each ROI signal *x*_*i*_(*t*) is binarized using a threshold *θ*_*i*_, where *i* ∈ {1, …, *N*}. The binarized value *σ*_*i*_(*t*) ∈ {−1, +1} is defined as:

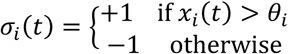

The full binarized vector at each time point *t* is:

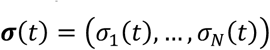

Each *σ*_*i*_(*t*) represents the binary activation state of the *i*-th ROI at time *t*, where +1 indicates “active” and −1 indicates “inactive”. Thus, the signal pattern at time *t* is given by the *N*-dimensional binary vector ***σ***(*t*), also referred to as an activity pattern or active pattern. For notational simplicity, the time index *t* may be omitted where appropriate. In total, there are 2^*N*^ possible activity patterns, representing all binary combinations of ROI states.

#### Fitting a pMEM

A pMEM is fitted to binarized time series data to estimate the energy associated with each observed signal pattern. For neuroimaging data, this allows modeling the distribution and dynamics of brain states based on observed frequencies of binary activation patterns. First, we compute the empirical distribution *P*_empirical_(***σ***) which represents the relative frequency of each activity pattern observed in the data (Figure 1D). Let ***σ***^(1)^, …, 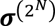 denote all possible binary activity patterns. Then, *P*_empirical_(***σ***) is simply the proportion of time points where pattern ***σ*** occurs, calculated by:

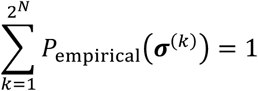

That is, the empirical frequency distribution *P*_empirical_(***σ***) is calculated by counting how many times each of the 2^*N*^ possible binary patterns appear in the binarized time series and dividing by the total number of time points. The empirical mean signal pattern of the *i*-th ROI is:

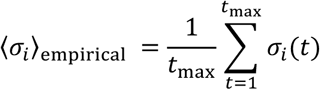

The empirical pairwise correlation between the *i*-th and *j*-th ROIs is:

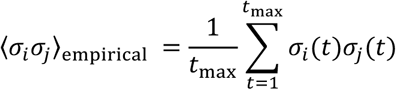

The pMEM estimates a probability distribution *P*(***σ***) that matches these first- and second-order moments while maximizing entropy. The result is the Boltzmann distribution:

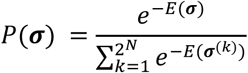

Equivalently, the distribution can be expressed in terms of model parameters ***h*** and ***J*** as:

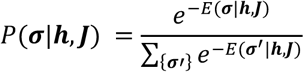

with the energy function:

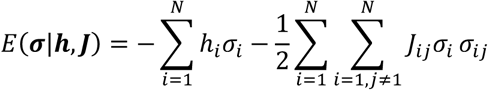

This formulation models the energy of each pattern based on ROI-wise signal tendencies (ℎ_*i*_) and pairwise interactions among ROIs (*J*_*ij*_).

In this study, the model parameters ***h*** and ***J*** were estimated by pseudo-likelihood maximization, following the implementation described by Ezaki et al. (2017). Full details of the fitting procedure, including gradient ascent optimization, are available in their original publication. Specifically, we used a Python implementation of the ELA toolbox developed by Oku (2023/2025) (https://github.com/okumakito/elapy).

#### Accuracy metrics for pMEM fitting

To evaluate the quality of the fitted pMEM, we used two metrics that quantify how well the model approximates the empirical distribution of signal patterns. The first indicator is defined as:

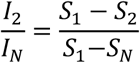

where *S*_*k*_ is the Shannon entropy of the MEM considering *k*-order correlations. It is calculated as:

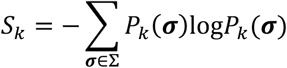

Here, *P*_1_(***σ***) corresponds to the independent MEM, *P*_2_(***σ***) to the pairwise MEM and *P*_*N*_(***σ***) to the empirical distribution observed in the data.

The second indicator is based on Kullback-Leibler (KL) divergence and is defined as:

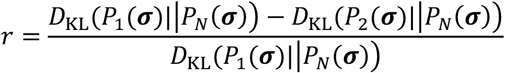

where the KL divergence is computed as:

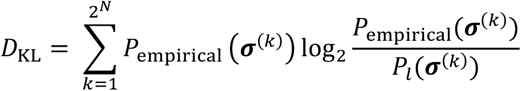

Here, *D*_KL_ quantifies the dissimilarity between the empirical distribution *P*_empirical_(***σ***) and the distribution of signal patterns estimated by the *l*-th order MEM *P*_*l*_(***σ***) (*l* = 1, 2).

In this study, the arithmetic mean of above two metrics was used as the quality measure of a pMEM fitting.

#### Personalization of pMEM via temperature optimization

In standard ELA, individual data is concatenated into a group-level dataset to estimate a single set of ***h*** and ***J*** parameters representing group characteristics. Although it is theoretically possible to fit pMEM to individual data, this approach is rarely adopted in practice because it requires a large amount of data to achieve stable estimation. For example, in typical fMRI datasets, the number of time points per individual is often insufficient to reliably estimate all parameters of the model. Therefore, pMEM is typically estimated using concatenated group-level data for robustness and accuracy. However, when individual variability needs consideration, personalization of the pMEM becomes essential. Ruffini et al. (2023) proposed a two-step pMEM parameter optimization method in which the inverse temperature parameter *β*, which characterizes the Boltzmann distribution, is optimized for each individual data while keeping the group-derived ***h*** and ***J*** parameters fixed. This approach allows group-level trends to be captured by ***h*** and ***J***, while individual differences are partly parameterized by *β* within this personalization scheme. Here, we use *β* as a practical personalization parameter without asserting it as the sole source of inter-individual variability.

Here, the distribution for the individual is defined as:

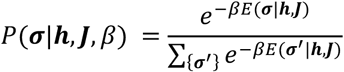

The optimization of *β* is performed by maximizing the approximate log-likelihood using a gradient ascent algorithm. The update rule is:

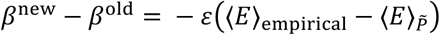

Where *ε* is the step size of the gradient ascent, 〈*E*〉_empirical_ is the average energy computed from the data, and 〈*E*〉_*P̃*_ is the model-based expected energy. These are derived from the log-likelihood expression:

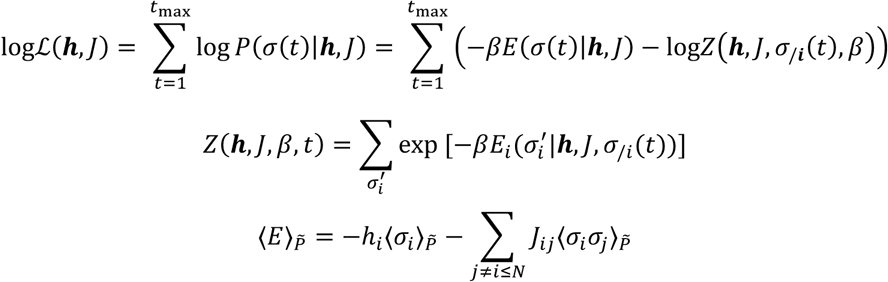

This procedure enables the personalization of the energy landscape while preserving group-level structure. The implementation in this study closely followed the approach described by Ruffini et al. (2023) and all equations and optimization steps are reproduced here for clarity and reproducibility.

#### Energy landscape and local minimum

By fitting a pMEM to the observed time series, the energy associated with any given pattern ***σ*** can be estimated. Since each pattern ***σ*** lies in a high-dimensional binary space defined over *N* ROIs, its neighboring patterns are those that differ from ***σ*** by exactly one bit (i.e., Hamming distance = 1). For example, consider ***σ*** = (1, 1, −1, 1) with *N* = 4. This pattern has four neighbors obtained by flipping each element. These *N* neighbors form vertices of an *N*-dimensional hypercube surrounding ***σ***. This structure allows us to interpret the temporal dynamics as transitions among vertices within this energy-defined space (that is, energy landscape).

Under the Boltzmann distribution, states with lower energy tend to appear more frequently, making local minima in the energy landscape particularly important. In ELA, such local minima are interpreted as representative signal patterns (states). The dynamics between these stable patterns, that is, how the system transitions from one to another, provide insights into the system structure. Moreover, the characteristics of the constructed energy landscape can be visualized using a contour plot, which highlights the energy barriers between local minima, as shown in Figure 1E.

### Automated ROI selection using a genetic algorithm

We introduce a mechanism to automatically determine the optimal subset of ROIs within the aforementioned ELA framework. In this study, we formulate ROI selection as a combinatorial optimization problem. To provide a representative example of how this optimization can be implemented, the goal is set to select a subset of ROIs that maximizes the quality of the pMEM fitting while capturing individual variability.

A critical aspect of combinatorial optimization is defining the objective function, which quantifies how well a candidate solution satisfies the desired properties. In this work, we define the objective function as the sum of:

- The fitting accuracy of the pMEM is represented by the similarity between the model and empirical distributions.
- The variance of the personalized inverse temperature parameter (*β*) across all participants, derived from the individualized Boltzmann distribution.

Here, the first term ensures the sufficient accuracy of pMEM fitting with the chosen ROIs, and the second is incorporated as a heuristic to maintain sensitivity to inter-individual variation in the energy landscape, preventing the selection of ROIs that merely maximize the fit to the population mean at the expense of inter-individual diversity. By using *β* as a parameter to account for these differences, we aim to identify ROI subsets that may better capture the individual dynamical characteristics of each participant’s brain state.

Let ***x*** ∈ {0,1}^*d*^ be a binary vector encoding the selection of *d* ROIs available, where *x*_*k*_ = 1 if the *k*-th ROI is selected and zero otherwise, and *N* ≤ *d* is the total number of ROIs to select. Then, the optimization problem is formulated as:

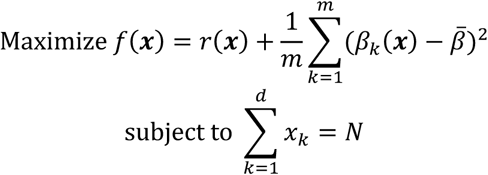

Here, *r*(***x***) denotes a pMEM fitting accuracy when ROIs specified by ***x*** are used, and *β*_*k*_(***x***) is the optimized inverse temperature for the *k*-th participant using the selected ROI subset ***x***. The mean value of *β*_*k*_(***x***) for *m* participants is denoted by 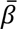. The constraint ensures that exactly *N* ROIs are selected from the *d* available candidates.

This objective function can be flexibly determined based on the analysis goals. For example, if the analyst expects the pMEM model parameters to correlate with data features, such correlations can be included as an additional term in the objective function. The objective function defined above serves as a representative instance to demonstrate the framework’s utility; however, the framework itself is independent to the specific choice of metrics, allowing users to incorporate various criteria depending on their research questions. By optimizing this function with a GA, the proposed framework enables data-driven selection of ROI subsets.

Regarding the search scope, the optimization process in ELA/GAopt performs a global search to identify a single, optimal ROI subset that is common across all participants within a study group. In this study, the optimization does not involve subject-level ROI selection; instead, it evaluates the fitness of candidate ROI sets based on the pMEM fitting accuracy and inter-individual variability, both of which are derived from the same cohort of participants. At each evaluation step of the GA’s objective function, the inverse temperature parameter *β* is individually optimized for every participant using the gradient ascent algorithm detailed in the previous section (“Personalization of pMEM via temperature optimization”). These optimized *β* parameters, derived for all participants, are then used to calculate the inter-individual variability term in the objective function.

For the implementation of a GA, we used a standard evolutionary framework based on Holland’s original design (Baeck et al., 2018), using the Deap library (F.-A. Fortin et al., 2012). We employed tournament selection, two-point crossover, and bit-flip mutation. To ensure the feasibility of the solutions, we incorporated a repair strategy called Lamarckian repair (Ishibuchi et al., 2005), which iteratively modifies infeasible solutions until they satisfy the specified constraints. Specifically, if the number of selected ROIs exceeds the prescribed limit, a subset of ROIs with a value of 1 (i.e., indicating selection) is randomly chosen and flipped to 0. The GA iteratively explored the optimal combinations of the ROIs for ELA. The full implementation details and parameter settings are provided in Algorithm 1 and Table 1, respectively.

#### Algorithm 1.

Procedure of a genetic algorithm.

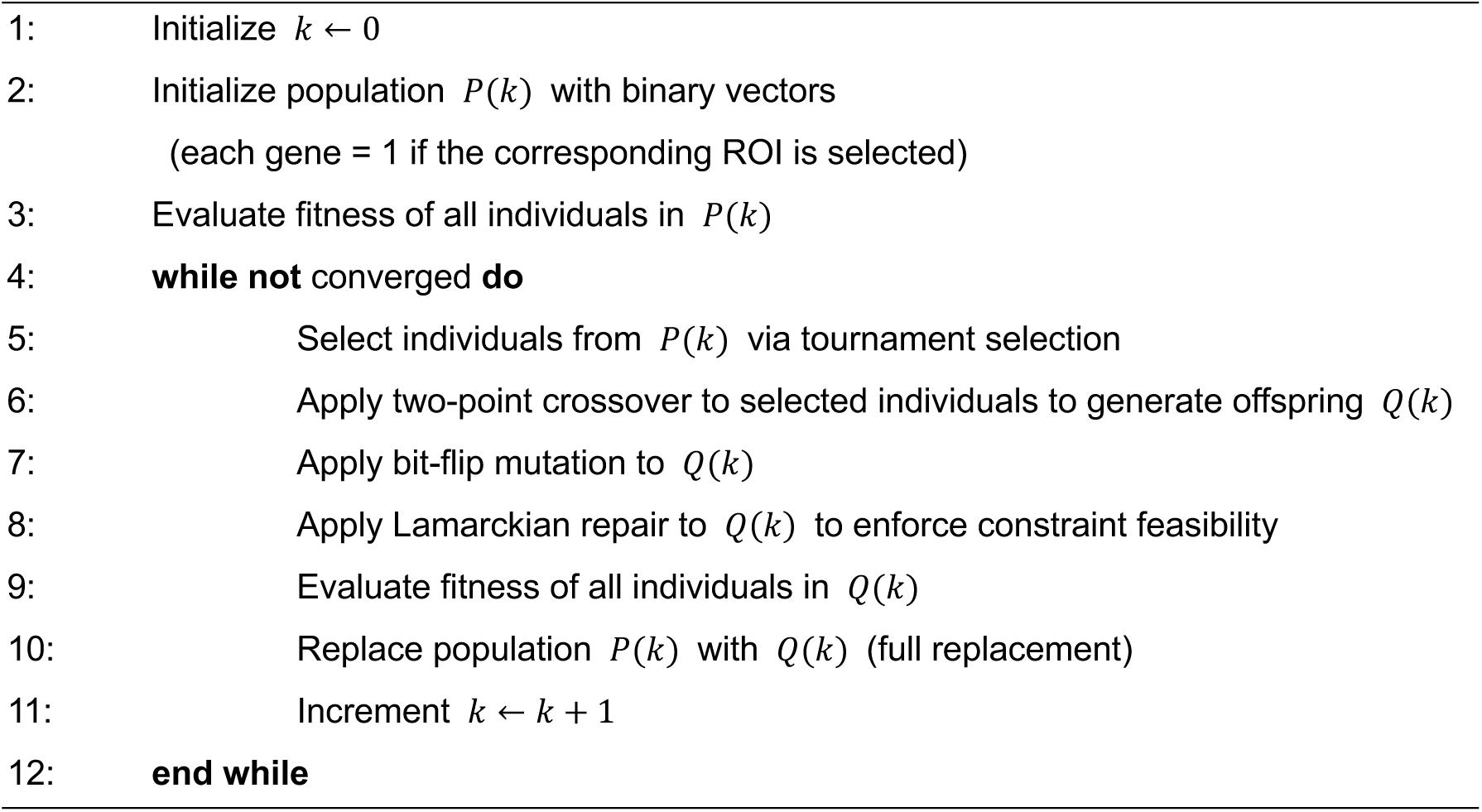

**Table 1.** Parameters of GA.

| GA Parameter | Value |
| --- | --- |
| Population size | 100 |
| Gene length | 160 for Scenario I (Creativity dataset),<br>264 for Scenario I (HCP-YA dataset) and Scenarios 2-4 |
| Number of generations | 1000 |
| Crossover method | Two-point crossover |
| Crossover rate | 1.0 |
| Mutation rate | 1/ Gene length |
| Selection method | Tournament selection (tournament size: 3) |
| Trial | 100 |

The source code for our proposed ELA/GAopt, along with example data and step-by-step usage instructions, is publicly available on GitHub at https://github.com/MIS-Lab-Doshisha/ela-gaopt. The repository includes a README file that details the installation process, data formatting, and execution of the pipeline. This facilitates easy adoption by new users.

### Datasets

To evaluate the performance and generalizability of the proposed ELA/GAopt framework, we applied it to three independent resting-state fMRI datasets. In the following sections, we overview the datasets used, including their preprocessing methods, as well as the analysis scenarios that utilize these datasets.

#### Creativity Dataset

We employed a publicly available dataset related to creativity, hosted on OpenNeuro (dataset ID: ds002330) (Sunavsky & Poppenk, 2020). A total of 66 healthy adults completed a neuroimaging session and four creativity assessments: the Abbreviated Torrance Test for Adults (ATTA) in verbal and visual formats, the Creative Behavior Inventory (CBI), and the Creative Achievement Questionnaire (CAQ). One participant did not complete the ATTA due to logistical reasons. The sample consisted of 29 males, 36 females, and one non-binary individual, with a mean age of 26.6 years (SD = 4.3).

MRI data included T1- and T2-weighted structural images, resting-state fMR images, and diffusion-weighted images (DWIs). After excluding participants with missing ATTA scores or insufficient data quality, the final analytic sample included 61 participants. These were stratified into a discovery set (*n* = 31) and a test set (*n* = 30) based on their ATTA verbal scores. The descriptive characteristics of two subsets are provided in Table 2.

**Table 2.** Basic descriptive characteristics of discovery and test sets.

|  | Data validation |  |
| --- | --- | --- |
|  | Discovery | Test |
| N | 31 | 30 |
| CBI score | 0.904 ± 0.509 | 0.941 ± 0.541 |
| ATTA verbal score | 16.300 ± 6.415 | 14.516 ± 6.597 |
| ATTA visual score | 29.017 ± 12.058 | 29.403 ± 10.139 |
| CAQ score | 4.172 ± 1.411 | 3.873 ± 1.769 |

#### ABIDE II Dataset

We used data from the Autism Brain Imaging Data Exchange II (ABIDE II), which provides a large multi-site resting-state fMRI and anatomical dataset for investigating ASD. Data from 17 sites were included after excluding two sites due to preprocessing incompatibility. The original dataset included 1,114 participants (ASD: 521, CTL: 593); 81 participants were removed due to preprocessing issues, resulting in a final sample of 1,033 participants.

Each remaining participant provided at least one resting-state fMRI scan and a high-resolution T1-weighted structural image. Basic demographic and cognitive data (e.g., age, sex, IQ) were available for all individuals. Detailed acquisition protocols and site-specific procedures are described in the related publications (Di Martino et al., 2014, 2017).

To balance site-specific variance, we grouped the dataset into three cohorts of approximately 300 participants each (A, C, and D) and an additional cohort (B) of around 100 participants used for independent validation. The distribution of participants across sites and cohorts is provided in Table 3.

**Table 3.** Distribution of creativity indicators in each cohort of ABIDE II Dataset.

| Cohort | Sites | Total N | N: ASD | N: Control |
| --- | --- | --- | --- | --- |
| A | NYUI, ONRC, SDSU, TCD,<br>UCD, UCLA | 293 | 157 | 136 |
| B | OHSU, USM | 126 | 54 | 72 |
| C | IU, KKI, KUL, NYU2 | 306 | 131 | 175 |
| D | BNI, EMC, ETH, GU, IP | 308 | 140 | 168 |

#### HCP-YA Dataset

We also utilized the Human Connectome Project Young Adult (HCP-YA) 2025 release dataset as a large-scale benchmark for healthy participants (Feinberg et al., 2010; Moeller et al., 2010; Setsompop et al., 2012; Van Essen et al., 2013; Xu et al., n.d.). The HCP-YA dataset provides a comprehensive investigation of brain function and structure in 1,113 healthy young adults aged 22–35 years. After excluding participants with missing data or insufficient image quality, a final analytic sample of 1,009 individuals was retained. From this sample, we extracted two independent subsets with balanced age distributions: a discovery set (*n* = 180) and a test set (*n* = 90). The discovery set had a mean age of 27.17 years (SD = 3.30), while the test set had a mean age of 27.54 years (SD = 3.77). A Welch’s t-test confirmed that there were no statistically significant age differences between these two subsets (*p* = 0.42).

#### Data preprocessing

Three independent datasets, the Creativity dataset, the ABIDE II dataset and HCP-YA dataset, were preprocessed in the following steps. It should be noted that the ABIDE II and HCP-YA data were originally provided in non-BIDS format and were converted to BIDS format using the procedure outlined by Ran et al. (https://github.com/thebrisklab/ABIDE_Preprocessing) (Brain Research in Imaging Statistics Kit, 2022/2024) and Suyash et al (https://github.com/suyashdb/hcp2bids) (Suyash, 2016/2025). For the ABIDE II dataset, the first five volumes were excluded to remove non-equilibrium effects of magnetization. This exclusion step was not applied to the Creativity dataset due to its limited number of subjects and time points. For the HCP-YA dataset, no slice timing correction was performed following to the prior study (S. M. Smith et al., 2013). Functional MR images from both datasets were then preprocessed using fMRIPrep version 23.1.4, a robust and standardized preprocessing pipeline built on Nipype.

First, a reference volume was generated from each run to use as a reference for head motion correction. The head motion parameters (transformation matrix and six parameters corresponding to translations and rotations) for this reference image were then estimated before spatiotemporal filtering was performed. Slice timing correction was further performed to correct the temporal misalignment of each slice. The EPI image was then aligned with the T1w image in six degrees of freedom using boundary-based registration.

Several confounding time series were subsequently estimated from the preprocessed BOLD data, including framewise displacements (FD), DVARS, and global signals from the cerebrospinal fluid (CSF), white matter (WM), and whole-brain mask.

Further preprocessing was carried out using the Nilearn package in Python (*Nilearn*, n.d.). This included spatial smoothing with a Gaussian kernel (FWHM = 6 mm), high-pass filtering at 0.01 Hz, low-pass filtering at 0.1 Hz, and regressing out of confounding variables (“high-pass”, “motion”, “wmcsf”, “scrub”). The denoising strategy followed established procedures in prior studies (H.-T. Wang et al., 2023).

Additionally, we used different atlases for each dataset: the Dosenbach atlas for Creativity Dataset (*d* = 160) (Dosenbach et al., 2006, 2007, 2010; Fair et al., 2009; Fox et al., 2005), and the Power 264 atlas for HCP-YA and ABIDE II Dataset (*d* = 264) (Power et al., 2011), respectively. The BOLD time courses were extracted from the spherical mask with a 4 mm radius centered at each ROI coordinate, then averaged within the ROI. The average time courses were considered representative of brain activity in each ROI. Finally, the ROI-averaged activity data was then binarized using the threshold of the time-averaged value of each activity for the ELA input.

### Statistical analysis

Statistical analyses were conducted independently for each scenario, as they addressed distinct research questions using different datasets. False Discovery Rate (FDR) correction using the Benjamini-Hochberg procedure was applied to all multiple comparisons, with the significance level set at adjusted *p* < 0.05. Specifically, FDR correction was applied to: (1) the comparison of optimized vs. randomly selected ROI sets in Scenario 1 (permutation tests for objective function values and Hamming distances), (2) the binomial tests for ROI selection frequency across all scenarios, and (3) the tests for occurrence probabilities of local minimum patterns in Scenario 2 (corrected for the number of identified patterns).

#### Scenario 1: Validation of generalizability of ELA/GAopt using the Creativity dataset and HCP-YA dataset

The objective of Scenario 1 was to evaluate whether the proposed automated ROI selection method avoids overfitting and selection bias. To ensure a rigorous evaluation, the partition between the discovery and test datasets was maintained at the subject level and remained fixed throughout across all 100 optimization runs. The GA-based optimization was conducted exclusively on the discovery dataset; the test dataset was completely excluded from the optimization process and reserved solely for independent validation. We applied ELA/GAopt to the discovery dataset in 100 independent runs, each with a different initial GA population and a different random seed, resulting in 100 optimized ROI sets.

Following Ezaki et al. (2017), who demonstrated that the accuracy of pMEM fitting scales with *l*/2^*N*^ (i.e., the average number of visits per activity pattern), the number of ROIs in Scenario 1(Creativity dataset) was fixed at 10 (*N* = 10). They reported that each activity pattern should be visited approximately 5 and 16 times on average to achieve accuracies of 0.8 and 0.9, respectively. For the *l* = 9170 data points in this scenario, choosing *N* = 10 (2^10^ = 1024) yields *l*/2^*N*^ ≈ 9.0 visits per pattern, which lies between 5 and 16; thus, we expect an accuracy between 0.8 and 0.9. Moreover, to ensure the generalizability of the results, we performed the same analysis on the HCP-YA dataset (*n* = 270). This larger sample size allowed us to further validate our method by testing a higher-dimensional model with 15 ROIs (*N* = 15), while still maintaining sufficient data volume per pattern to ensure robust pMEM fitting.

With the resulting ROIs chosen by ELA/GAopt for 100 trials, a one-tailed binomial test was performed to assess whether each brain region was selected more frequently than expected by chance. False discovery rate (FDR) correction was performed for multiple comparisons. To quantify the effect size for these binomial comparisons, Cohen’s ℎ was calculated. This process ensured the detection of consistently selected ROIs for multiple GA optimization runs.

Furthermore, to quantify the stability of ROI selection across the 100 independent GA runs, we conducted a pairwise similarity analysis. Each optimized ROI set was represented as a binary vector **v** ∈ {0, 1}^*d*^, where *d* denotes the total number of candidate regions in the atlas; elements were set to 1 for selected ROIs and 0 otherwise. We then calculated pairwise overlaps across all possible combinations of the 100 optimized ROI sets (7^100^) = 4,950) using two metrics: the Hamming distance, which counts the number of differing bits between two vectors, and the Jaccard index, defined as the size of the intersection divided by the size of the union of the selected ROI sets. For comparison, these metrics were also computed for 100 randomly selected ROI sets. The distributions of these metrics were compared between the optimized and random groups using the Mann-Whitney *U* test, with statistical significance assessed at *p* < 0.05. Additionally, Cohen’s d was calculated to quantify the effect size of the observed differences in stability between the optimized and random ROI sets.

Next, each of the 100 optimized ROI sets was applied to the test dataset using standard ELA. In parallel, we generated 100 randomly selected ROI sets, each also consisting of 10 ROIs for Creativity data and 15 ROIs for HCP-YA data. Then we applied them to both the discovery and test datasets using the same ELA procedure. In total, this yielded two groups of ROI sets, optimized, and randomly selected, for comparative analysis.

We tested two hypotheses: 1) the optimized ROI sets provide better objective function values (i.e., the sum of pMEM fitting accuracy and inter-individual variance of the inverse temperature) than random sets, and this advantage generalizes to the test data, and 2) the local minimum patterns identified in the discovery dataset are reproducible in the test dataset.

For Hypothesis 1, we compared the objective function values of the optimized and random ROI sets on the test dataset using permutation testing. Specifically, the objective values from both sets were pooled and the group labels (optimized vs. random) were shuffled 10,000 times to generate a null distribution. For each permutation, the mean objective value for each group was computed. A *p*-value was then derived by comparing the actual difference in mean objective values between the optimized and random groups to this null distribution. Statistical significance was assessed at *p* < 0.05. In addition, effect size (Cohen’s *d*) was calculated to quantify the observed differences between optimized and random ROI sets.

For Hypothesis 2, we quantified the similarity of local minimum patterns between the discovery and test datasets. Each pattern among the 2^*N*^ possible signal patterns (for *N* ROIs) were assigned a unique ID. For each ELA run, we generated an ID vector indicating the presence (1) or absence (0) of each pattern as a local minimum in the estimated energy landscape. We then calculated the Hamming distance between ID vectors from the discovery and test datasets for each ROI set. A smaller Hamming distance indicates greater reproducibility of the local minimum patterns. The Hamming distances of the optimized and random ROI sets were compared using a permutation test (with 10,000 permutations), with significance assessed at *p* < 0.05. In addition, Cohen’s *d* was computed to quantify the effect size of the difference in the Hamming distance. All permutation tests were one-tailed.

#### Scenario 2: Group-level comparison of brain dynamics on the ABIDE II Dataset

Scenario 2 investigated whether the different brain dynamics of two separate groups were observed on the single energy landscape constructed using ELA/GAopt on the concatenated dataset. Specifically, we aimed to observe group-specific brain dynamics on the energy landscape estimated on the mixed dataset of ASD and CTL groups, in a fully data-driven manner using ELA/GAopt. To confirm this, we extract the local minimum patterns more frequently visited in each group.

To ensure a rigorous evaluation and avoid selection bias, the partition between the discovery and test datasets was maintained at the subject level and remained fixed across all optimization runs. The GA optimization was conducted exclusively on the discovery dataset (Cohort A: *n* = 293; 157 ASD and 136 CTL); the test dataset (Cohort B: *n* = 126; 54 ASD and 72 CTL) was completely excluded from the optimization process and reserved solely for independent validation. The number of selected ROIs for the ABIDE II dataset in Scenarios 2 was fixed at 13, following the same procedure as in Scenario 1. Regarding the search scope, ELA/GAopt performed 100 independent optimization runs on the entire discovery cohort to identify single, optimal ROI subsets common across all participants in the cohort. Then, we evaluated how frequently each ROI was selected across trials. To determine whether specific ROIs were selected more frequently than expected by chance, a binomial test was performed for each ROI. The test assessed whether the observed selection frequency exceeded the expected frequency under a null hypothesis of random selection. The resulting *p*-values were corrected for multiple comparisons using FDR correction, and statistically significant ROIs (FDR-adjusted *p* < 0.05, one-tailed) were identified. To quantify the magnitude of deviation from chance-level selection, Cohen’s ℎ was calculated as the effect size. Furthermore, to quantify the algorithmic stability, we evaluated the consistency of the optimized ROI sets using the pairwise Jaccard index and Hamming distance, following the same procedure described in Scenario 1. ROI sets identified as significant in the binomial tests were considered consistently important and served as the basis for further group comparisons.

To perform group-level comparisons of local minimum patterns, we selected representative ELA/GAopt trials in which all selected ROIs were among those identified as statistically significant in the above binomial tests. These representative trials then estimated energy landscapes, and the probability of visiting each local minimum was computed for all individuals.

To compare these probabilities between the ASD and CTL groups, we applied the Mann-Whitney *U* test for each identified local minimum (FDR-adjusted *p* < 0.05, two-tailed). To quantify group difference magnitude, we also calculated the effect size (*r*) based on the *Z*- statistic from the Mann-Whitney *U* test. To evaluate the robustness of observed group-level differences, the same analysis was repeated on an independent validation dataset (Cohort B), following the same ROI configuration and statistical testing procedures.

#### Scenarios 3 and 4: Estimation of ASD- or CTL-optimized energy landscapes on the ABIDE II Dataset

In Scenario 3, we investigated whether a set of ROIs optimized for the ASD group could yield distinctive brain state characteristics. The GA-based optimization was performed exclusively on the ASD group (Cohort C, *n* = 131); to prevent overfitting and selection bias, the CTL group (Cohort C, *n* = 175) from the same cohort was maintained as an independent test set and was completely excluded from the optimization process. The number of selected ROIs was fixed at 12, following the same procedure as in Scenario 1. As in Scenario 2, we assessed the selection frequency of each ROI across trials and identified consistently selected ROIs using a binomial test (FDR-adjusted *p* < 0.05, one-tailed). For these binomial tests, Cohen’s ℎ was calculated as the effect size.

The resulting ASD-optimized ROI set was then applied to an independent CTL dataset (175 participants from Cohort C) to evaluate whether the same ROI configuration would produce distinct energy landscape characteristics. Specifically, the number of local minima identified in the CTL group was compared to that in the ASD group using the Mann-Whitney *U* test. To quantify the magnitude of group differences, we also calculated the effect size (*r*) based on the *Z*-statistic from the Mann-Whitney *U* test (two-tailed). We hypothesized that if the ASD-optimized ROIs reflect ASD-specific brain state dynamics, applying them to CTL data would result in significantly different energy landscape structures—particularly in the number or configuration of local minima—between the two groups.

Scenario 4 mirrored this procedure for the CTL group. The GA optimization was conducted exclusively on the CTL participants (Cohort D, *n* = 168), while the ASD group (Cohort D, *n* = 140) served as an independent test set. The optimization involved 100 global runs on the CTL discovery cohort to identify single, optimal ROI subsets. ROI selection analysis and statistical testing were conducted in the same manner as in Scenario 3. We hypothesized that if the CTL-optimized ROIs captured control-specific dynamics, applying them to ASD data would similarly yield distinct energy landscape structures. Finally, to quantify the algorithmic stability of ROI selection, we calculated the pairwise Jaccard index and Hamming distance across the 100 runs for both groups, following the same procedure described in Scenario 1.

#### Computational environment

We used a computing workstation equipped with an Intel Core (TM) i9-14900K CPU, a 7.7 GHz 24-core processor, and an NVIDIA GeForce RTX 4080 16GB GPU for ELA/GAopt execution.

## Result

### Scenario 1

First, we summarize the results of Creativity dataset. Supplementary Figure 1 shows the convergence of the objective function with ELA/GAopt over 1000 generations for 100 runs with different starting populations. As the optimization problem was formulated as a maximization task, the objective function value increased with each generation. All runs converged to a range between 1.00 and 1.10.

Based on the 100 ELA/GAopt runs, a binomial test was conducted to assess whether each ROI was selected more frequently than expected by chance. Eighteen ROIs were statistically significant after FDR correction (*p* < 0.05), as listed in Figure 2 and Supplementary Table 1, which also includes the anatomical coordinates and associated functional networks for each ROI. Additionally, the distribution of selected ROIs across the 100 runs is presented in Supplementary Figure 2, which shows the total number of times each ROI was selected. Furthermore, evaluation of the stability of the entire ROI set across repeated runs are shown in Figure 3. The pairwise Hamming distances between optimized ROI sets (ROI selection vectors) were significantly smaller than those between random sets (Mann-Whitney *U* test, *p* < 0.0001, FDR-corrected, Cohen’s *d*: 3.57, Figure 3A). Additionally, the pairwise Jaccard index between optimized ROI sets was significantly higher than that of randomly selected sets (Mann-Whitney *U* test, *p* < 0.0001, FDR-corrected, Cohen’s *d*: 2.83, Figure 3B). These results demonstrated that ELA/GAopt exhibited high reproducibility in ROI selection.

**Figure 2.**
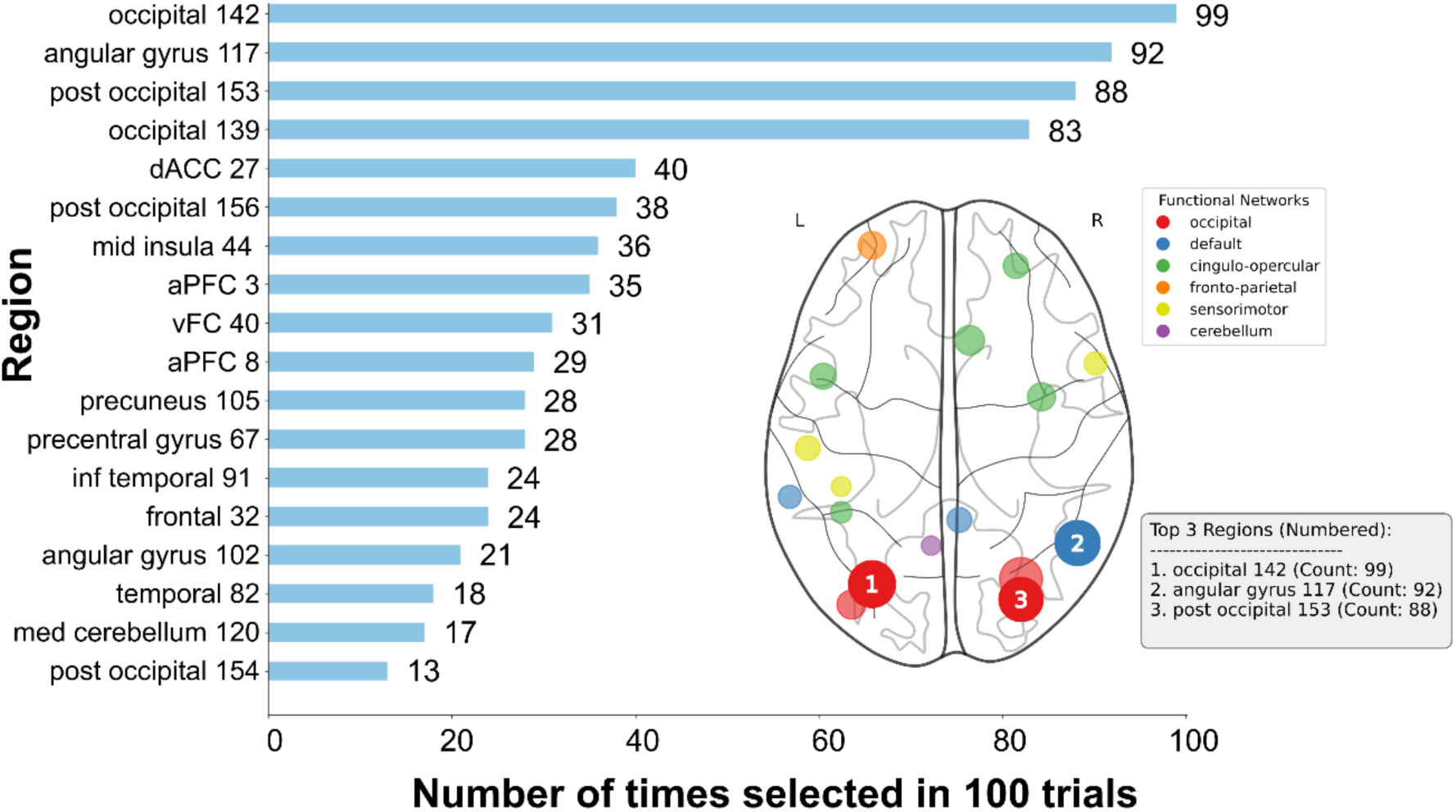
Selection frequency of each ROI across 100 runs of ELA/GAopt (Scenario 1, on Creativity Data). The bar graph illustrates the frequency with which each ROI was selected across 100 independent optimization runs. A binomial test was used to determine if the frequency of each ROI was higher than chance, and 18 ROIs were found to be statistically significant (***p*** < **0**. **0.5**, FDR-corrected) and indicated in the figure. The ROIs are arranged in order from most to least frequently selected. Their anatomical locations are also displayed on the glass brain plot. The circle size in the brain plot represents the frequency at which each ROI was chosen, with larger circles indicating higher frequencies.

**Figure 3.**
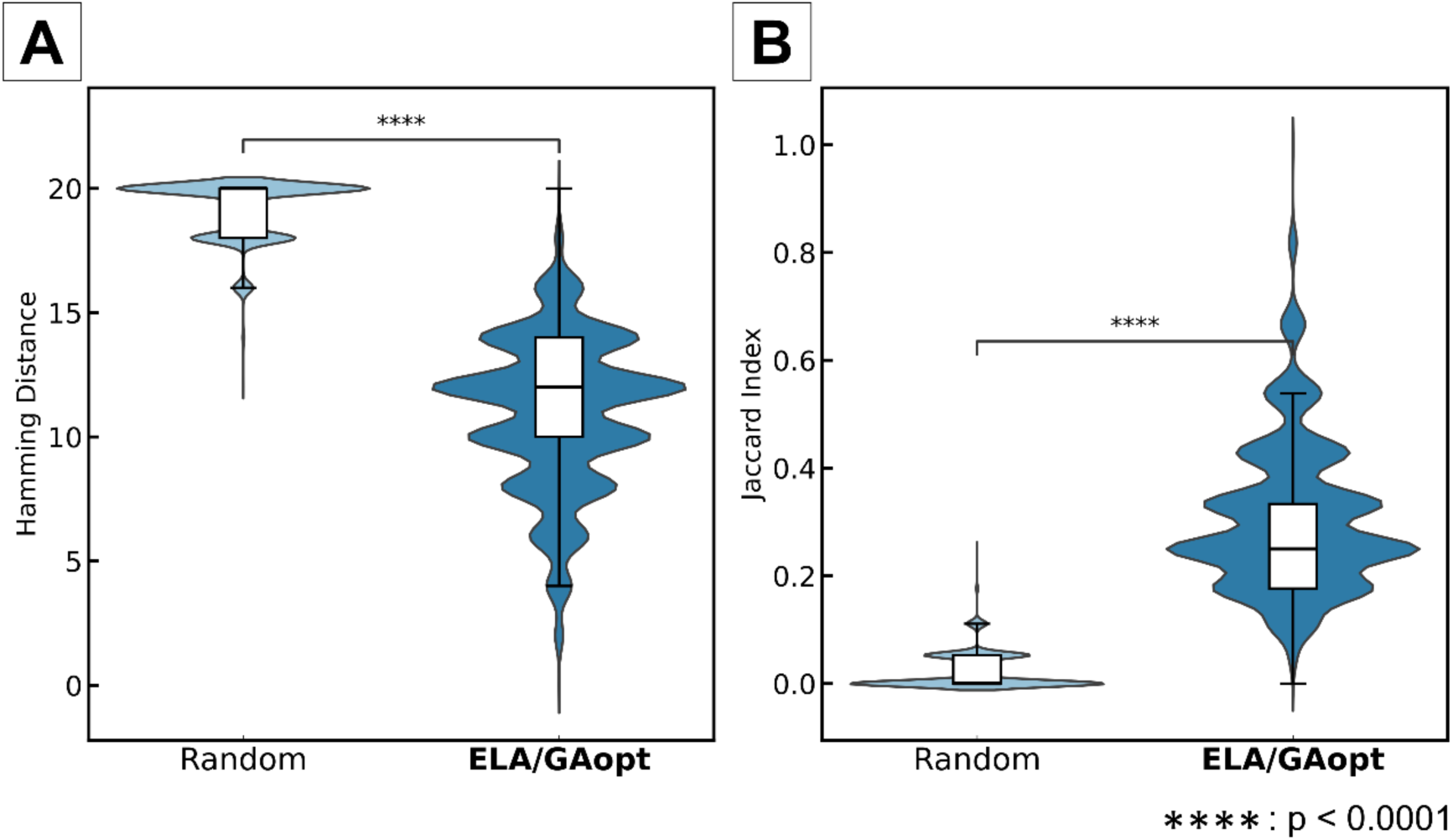
Stability evaluation of ROI selection using ELA/GAopt compared to random selection (Scenario 1, on Creativity data). (**A**) The pairwise Hamming distances between ROI sets (ROI selection vectors). The pairwise Hamming distances between optimized ROI sets were significantly smaller than those between random sets (Mann-Whitney U test, *p* < 0.0001, FDR-corrected, Cohen’s *d* = 3.57). (**B**) The Jaccard index between ROI sets (ROI selection vectors). The pairwise Jaccard index between optimized ROI sets was significantly higher than that between randomly selected sets (Mann-Whitney U test, *p* < 0.0001, FDR-corrected, Cohen’s *d* = 2.83).

To test the reproducibility of the local minimum patterns across datasets, we calculated the Hamming distance between ID vectors of the discovery and test datasets. Figure 4A and D show that the Hamming distances were significantly smaller for the optimized ROIs compared to the randomly selected ROIs (permutation test, 10,000 permutations, *p* = 0.0025, FDR-corrected, Cohen’s *d*: 0.408), indicating greater pattern stability with the optimized ROIs.

**Figure 4.**
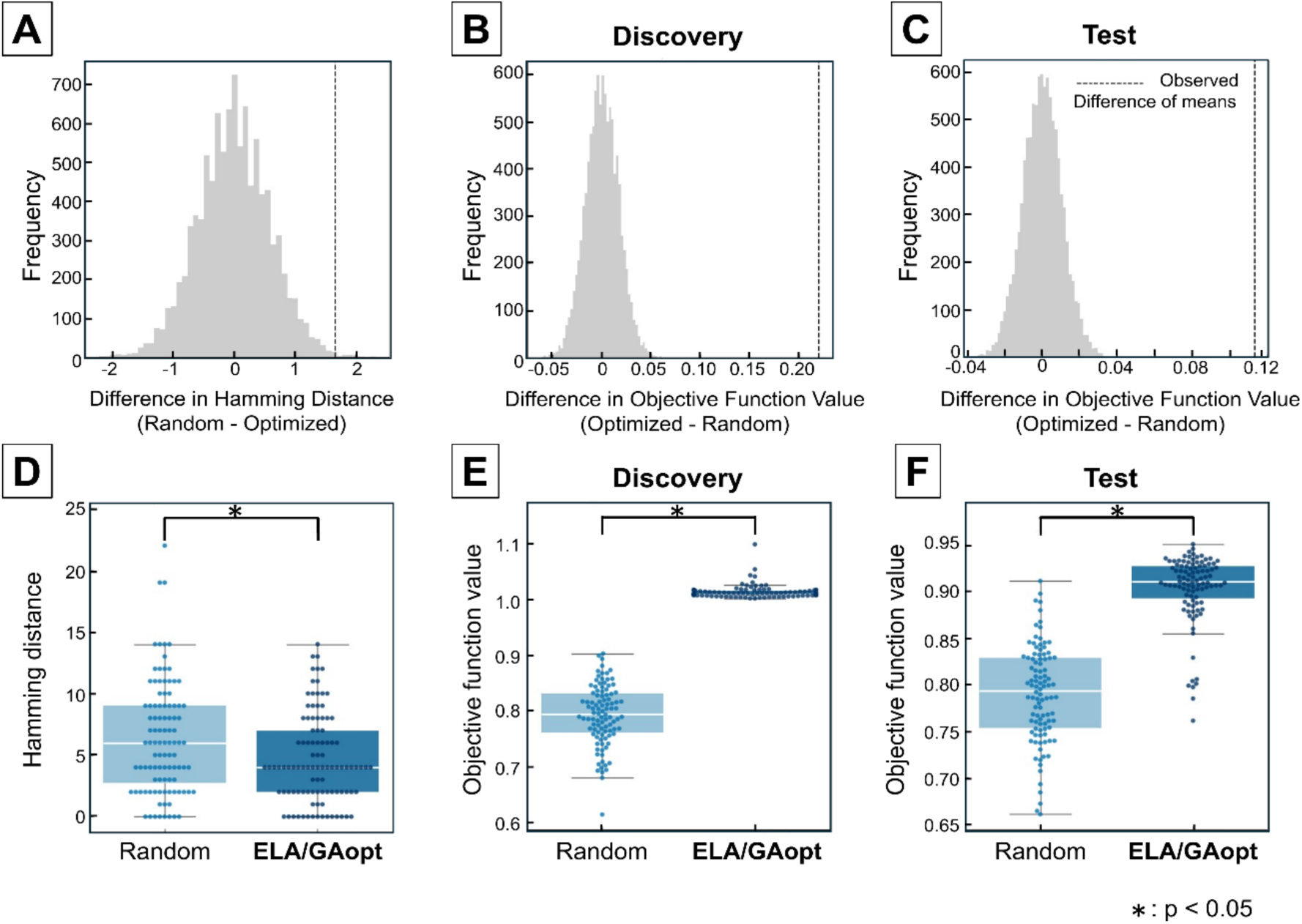
Reproducibility of local minimum patterns across discovery and test datasets (on Creativity Data). (**A** - **C**) Null distributions (10,000 permutations) of the difference in mean values between optimized and randomly selected ROI sets: (**A**) Hamming distance, (**B**) objective function value in the discovery set, and (**C**) objective function value in the test set. (**D**) The Hamming distances between local-minima state ID vectors of the discovery (*n* = 31) and test (*n* = 30) datasets. The Hamming distances were significantly smaller for the optimized ROI sets compared to the randomly selected ROIs (permutation test, *p* = 0.0025, FDR- corrected, Cohen’s *d* = 0.408). (**E**) The objective function values in the discovery data for the optimized ROI sets were significantly higher than those of the randomly selected ROI sets (permutation test, *p* < 0.001, FDR-corrected, Cohen’s *d* = 5.63). (**F**) The significant advantage of optimized ROI sets also generalized to the test set when compared to randomly selected ROI sets (permutation test, *p* < 0.001, FDR-corrected, Cohen’s *d* = 2.51). We generated 100 optimized ROI sets and 100 randomly selected ROI sets.

Finally, Figure 4B, C, E, and F summarize the comparison of the objective function values in both the discovery and test sets. Permutation testing revealed that the values for the optimized ROI sets were significantly higher than those of the randomly selected sets in the discovery set (*p* < 0.001, FDR-corrected, Cohen’s *d*: 5.63). Crucially, this significant advantage also generalized to the independent test set (*p* < 0.001, FDR-corrected, Cohen’s *d*:2.51), confirming the generalizability and effectiveness of the proposed ROI selection framework.

Next, results using the HCP-YA dataset are described. Supplementary Figure 4 shows the convergence of the objective function over 1000 generations using ELA/GAopt across 100 runs with different initial populations. All runs converged within the range of 0.96 to 0.98 objective function value.

With the 100 ELA/GAopt runs, binomial tests were performed to evaluate whether each ROI was selected more frequently than expected by chance. After FDR correction (*p* < 0.05), 20 ROIs were statistically significant as shown in Figure 5. Supplementary Table 2 lists the anatomical coordinates and associated functional networks for each ROI. Furthermore, the distribution of selected ROIs across 100 runs is shown in Supplementary Figure 5, where the total number of selections for each ROI can be confirmed.

**Figure 5.**
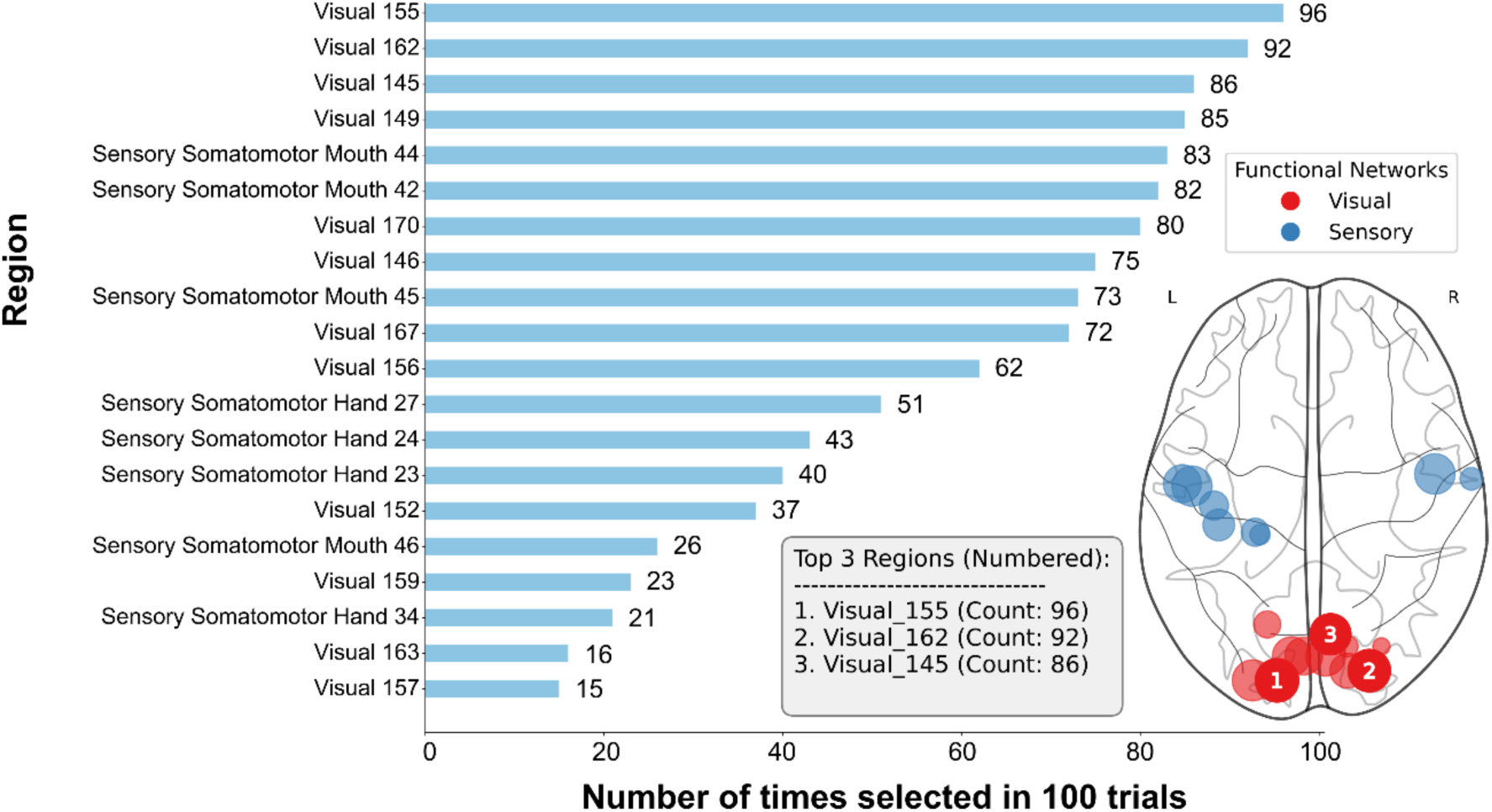
Selection frequency of each ROI across 100 runs of ELA/GAopt (Scenario 1, on HCP-YA Data). The bar graph illustrates the frequency with which each ROI was selected in 100 independent optimization runs. A binomial test was used to determine if the selection frequency of each ROI was higher than chance, and 20 ROIs were found to be statistically significant (*p* < 0.05, FDR-corrected), as shown in the figure. The ROIs are arranged in order from most to least frequently selected. Their anatomical locations are also displayed on the glass brain plot. The circle size in the brain plot represents the frequency of each ROI’s selection, with larger circles indicating higher frequencies.

The stability of the optimized ROI set across iterative runs were also evaluated using HCP-YA dataset. As shown in Figure 6, the pairwise Hamming distances between ROI selection vectors were significantly smaller than those between random sets (Mann-Whitney U test, *p* < 0.0001, FDR-corrected, Cohen’s *d*: 5.81, Figure 6A). Additionally, the pairwise Jaccard index between optimized ROI sets was significantly higher than that of randomly selected sets (Mann-Whitney *U* test, *p* < 0.0001, FDR-corrected, Cohen’s *d*: 4.71, Figure 6B). These results confirmed that ELA/GAopt exhibited high reproducibility in ROI selection, also in HCP-YA dataset.

**Figure 6.**
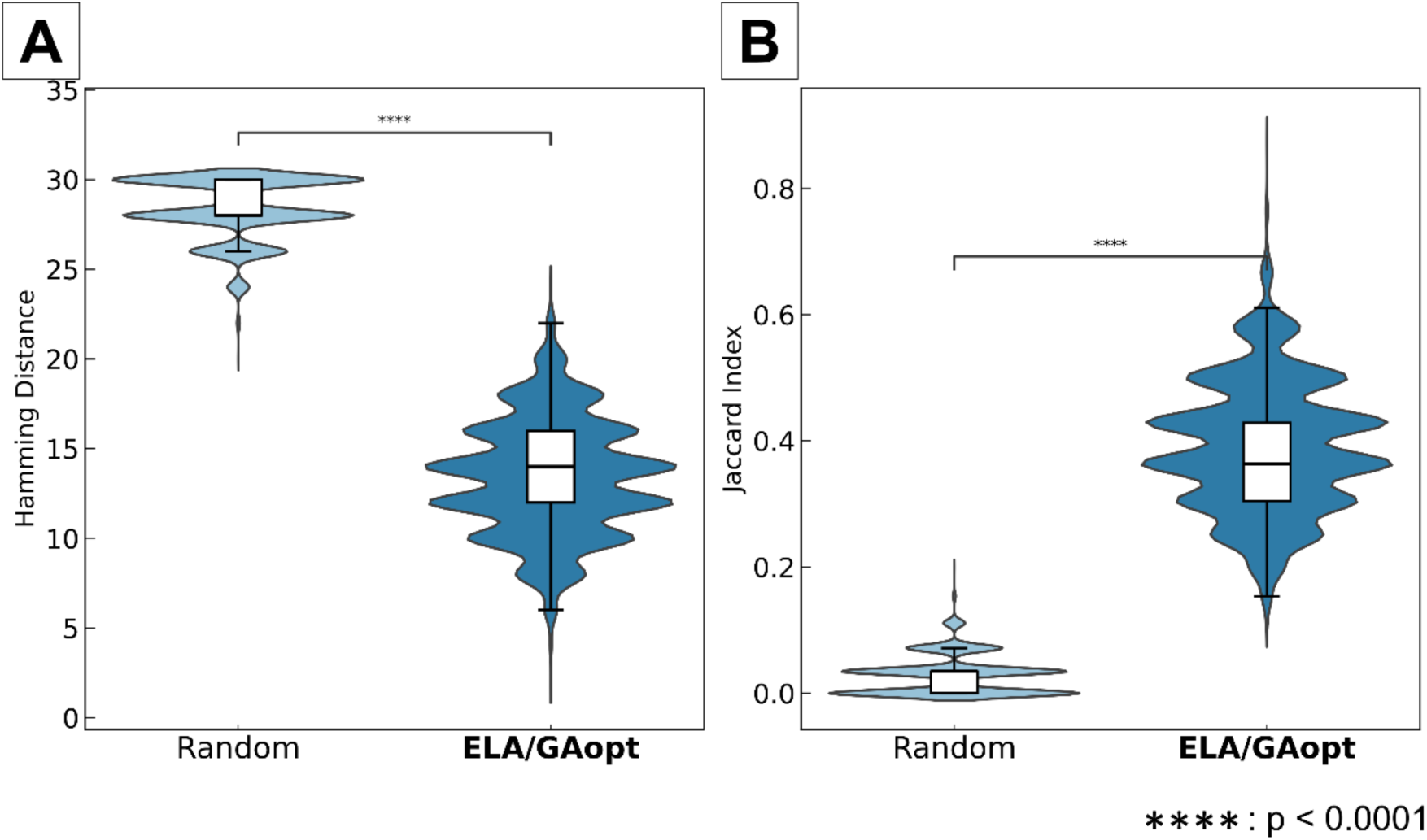
Stability evaluation of ROI selection using ELA/GAopt compared to random selection (Scenario 1, on HCP-YA data). (**A**) The pairwise Hamming distances between ROI sets (ROI selection vectors). The pairwise Hamming distances between optimized ROI sets were significantly smaller than those between random sets (Mann-Whitney U test, *p* < 0.0001, FDR-corrected, Cohen’s *d* = 5.81). (**B**) The Jaccard index between ROI sets (ROI selection vectors). The pairwise Jaccard index between optimized ROI sets was significantly higher than that between randomly selected sets (Mann-Whitney U test, *p* < 0.0001, FDR-corrected, Cohen’s *d* = 4.71).

Regarding objective function values, the optimized set showed significantly higher values than the random set in both the discovery dataset (*p* < 0.01, FDR-corrected, Cohen’s *d*: 6.22, Figure 7B, E) and the independent test dataset (*p* < 0.01, FDR-corrected, Cohen’s ***d***: 6.95, Figure 7C, F). In contrast, for pattern reproducibility quantified by the Hamming distance between local-minima states, we observed a trend opposite to that in the Creativity dataset. A one-tailed permutation test in the hypothesized direction (ELA/GAopt < Random) did not support a reduction in Hamming distance for the optimized ROI sets relative to the random ROI sets (*p* = 0.99, Figure 7A, D), indicating an effect opposite to the expected direction. We discuss possible reasons for this discrepancy from the Creativity dataset in the Discussion. As a post-hoc analysis, the optimized ROI sets contained a significantly larger number of identified local-minima states than the random ROI sets (discovery data: *p* = 0.0085, Cohen’s *d* = 0.34; test data: *p* = 0.0001, Cohen’s *d* = 0.54). After controlling for this difference in state count using a normalized comparison, we still observed no evidence that optimization reduced Hamming distance (permutation test, *p* = 0.99). Furthermore, the number of consistent local-minimum states did not differ between optimized and random ROI sets (permutation test, *p* = 0.43).

**Figure 7.**
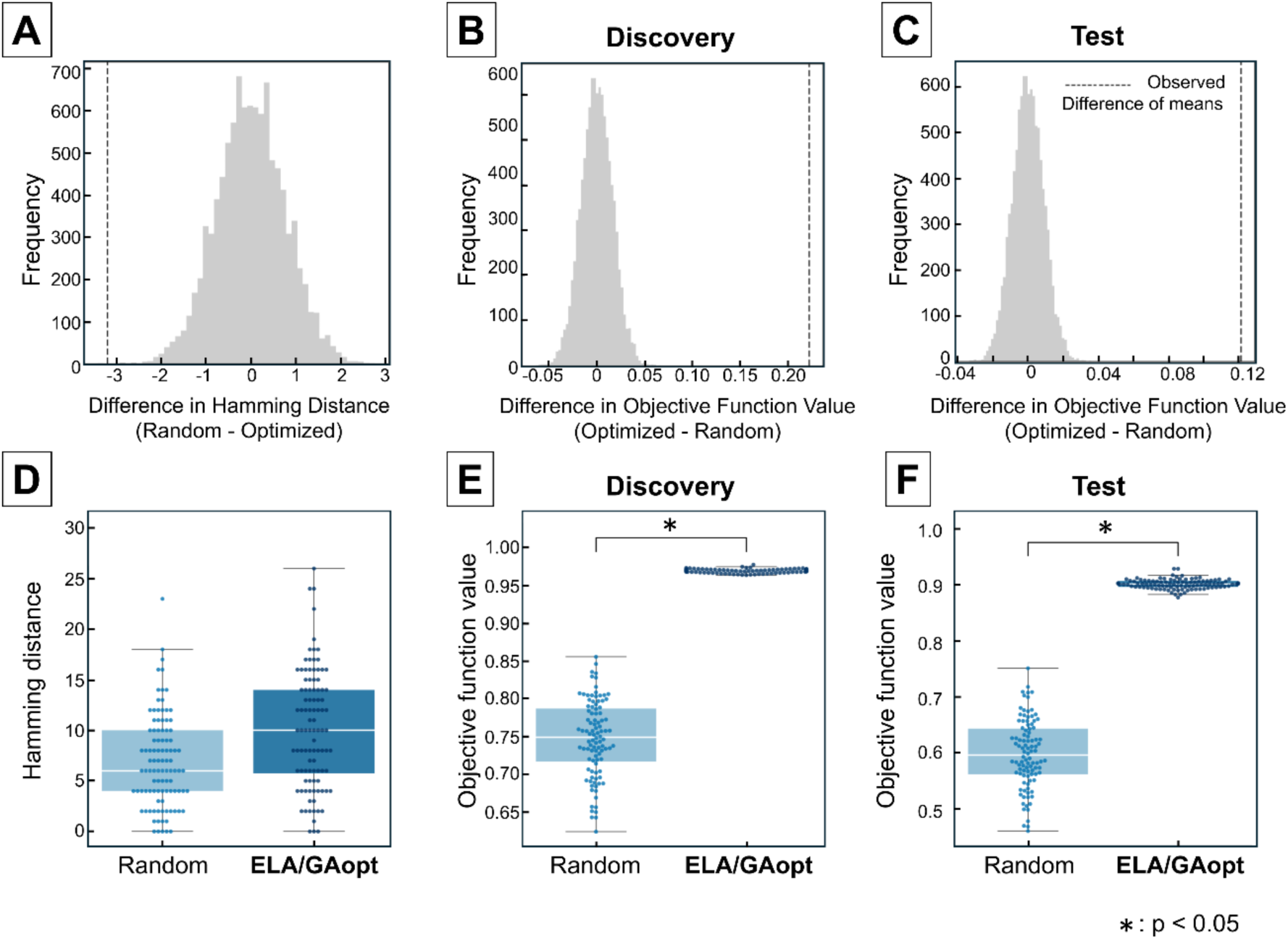
Reproducibility of local minimum patterns across discovery and test datasets (on HCP-YA Data). (**A** - **C**) Null distributions (10,000 permutations) of the difference in mean values between optimized and randomly selected ROI sets: (**A**) Hamming distance, (**B**) objective function value in the discovery set, and (**C**) objective function value in the test set. The vertical dashed line indicates the observed mean difference. (**D**) The Hamming distances between local-minima state ID vectors of the discovery (*n* = 180) and test (*n* = 90) datasets. The optimized ROI sets did not yield a significantly smaller Hamming distance compared to the randomly selected ROI sets (one-tailed permutation test, hypothesized direction: ELA/GAopt < Random, *p* = 0.99). (**E**) The objective function values in the discovery data for the optimized ROI sets were significantly higher than those of the randomly selected ROI sets (permutation test, *p* < 0.001, FDR-corrected, Cohen’s *d* = 6.22). (**F**) The significant advantage of optimized ROI sets also generalized to the test set when compared to randomly selected ROI sets (permutation test, *p* < 0.001, FDR-corrected, Cohen’s *d* = 6.95). We generated 100 optimized ROI sets and 100 randomly selected ROI sets.

### Scenario 2

Supplementary Figure 6 illustrates the convergence of the objective function over 1000 generations during ELA/GAopt, conducted in 100 independent runs with different initial populations. As with Scenario 1, the optimization process consistently improved the objective function value, with most runs converging by the end of the generation window.

Based on the ROI selection results from the 100 runs, we performed a statistical analysis using a binomial test to determine whether each brain region was selected more frequently than expected by chance. The resulting ***p***-values were adjusted using the FDR correction to account for multiple comparisons. Brain regions identified as significant are presented in Figure 8 and Supplementary Table 3. Furthermore, stability analysis confirmed the robustness of ROI selection in Scenario 2 (Figure 9). The optimized ROI set exhibited significantly lower Hamming distance between ROI selection vector compared to random selection (Mann-Whitney *U* test, *p* < 0.0001, FDR-corrected, Cohen’s *d*: 4.90, Figure 9A) and a higher Jaccard index (Mann-Whitney *U* test, *p* < 0.0001, FDR-corrected, Cohen’s *d*: 4.32, Figure 9B).

**Figure 8.**
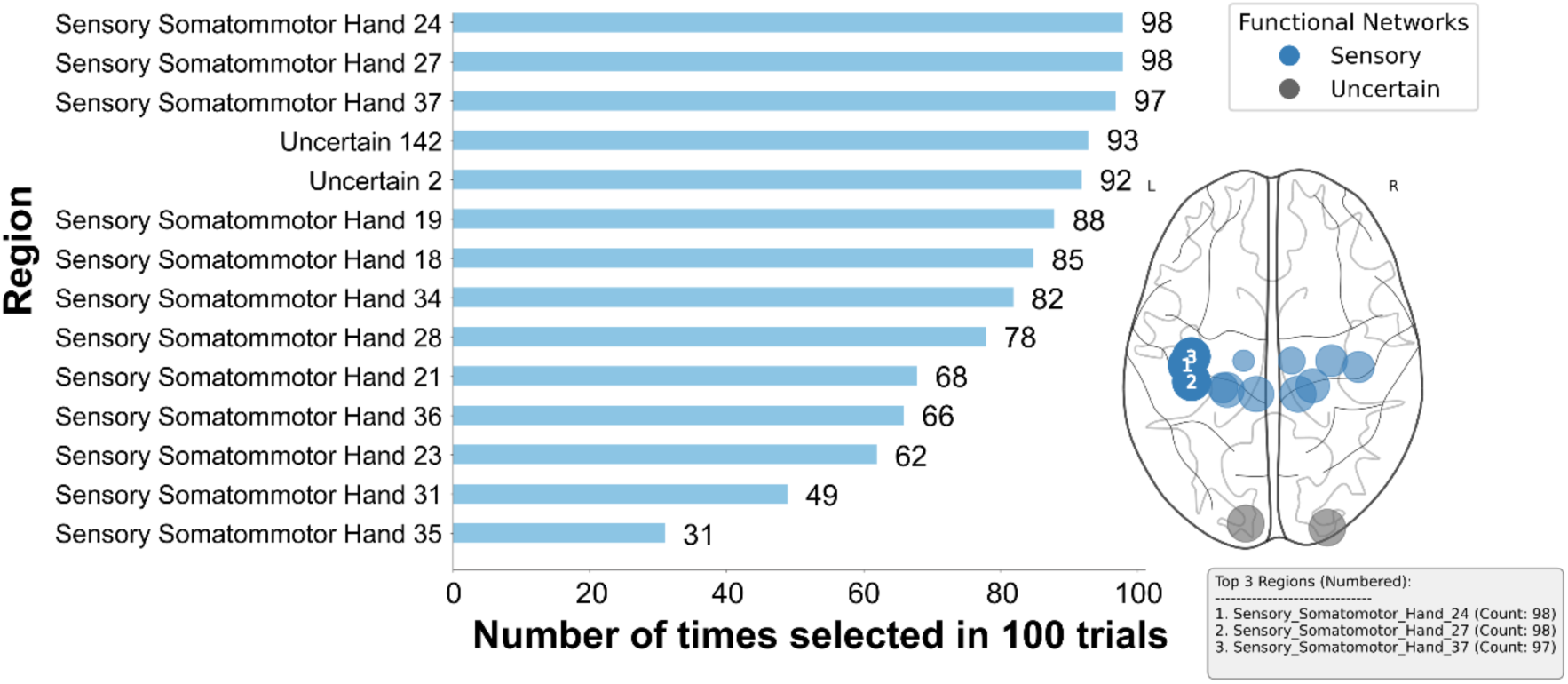
Selection frequency of each ROI across 100 runs of ELA/GAopt (Scenario 2). The bar graph illustrates the frequency with which each ROI was selected in 100 independent optimization runs. A binomial test was used to determine if the selection frequency of each ROI was higher than chance, and 14 ROIs were found to be statistically significant (*p* < 0.05, FDR-corrected), as shown in the figure. The ROIs are arranged in order from most to least frequently selected. Their anatomical locations are also displayed on the glass brain plot. The circle size in the brain plot represents the frequency of each ROI’s selection, with larger circles indicating higher frequencies.

**Figure 9.**
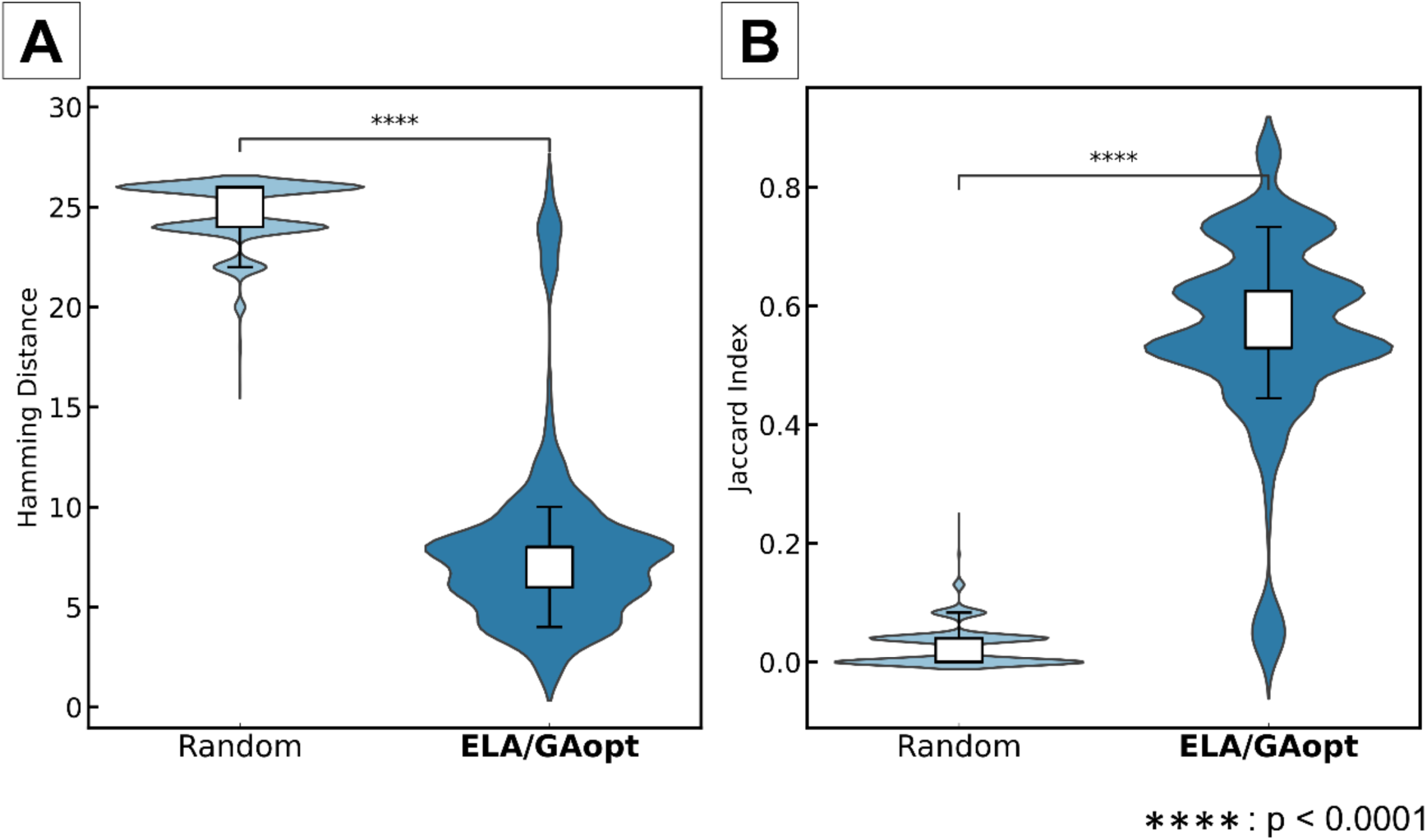
Stability evaluation of ROI selection using ELA/GAopt compared to random selection (Scenario 2). (**A**) The pairwise Hamming distances between ROI sets (ROI selection vectors). The pairwise Hamming distances between optimized ROI sets were significantly smaller than those between random sets (Mann-Whitney U test, *p* < 0.0001, FDR-corrected, Cohen’s *d* = 4.90). (**B**) The Jaccard index between ROI sets (ROI selection vectors). The pairwise Jaccard index between optimized ROI sets was significantly higher than that between randomly selected sets (Mann-Whitney U test, *p* < 0.0001, FDR-corrected, Cohen’s *d* = 4.32).

Additionally, the overall distribution of ROI selections across the 100 trials is presented in Supplementary Figure 7. The selection frequency indicates how often ROIs from each functional network were selected during the automated ROI selection process in ELA/GAopt.

To compare the probability of visiting each local minimum between the ASD and CTL groups, a representative trial was selected from the 100 ELA/GAopt runs. Since the objective function values were sufficiently high across all trials, they were not suitable as selection criteria. Instead, we focused on two trials where all selected ROIs were identified as being significantly more frequently selected than expected by chance, based on the binomial test with FDR correction (*p* < 0.05).

In the first representative trial, 13 ROIs were selected (Figure 10A), and 12 distinct local minimum states were identified in Cohort A (Figure 10A, left panel). To assess group differences in the probability of visiting each of the four most prominent local minima (as listed in Table 4), Mann-Whitney *U* tests were conducted, followed by FDR correction. Significant between-group differences were observed (FDR-corrected *p* < 0.05).

**Figure 10.**
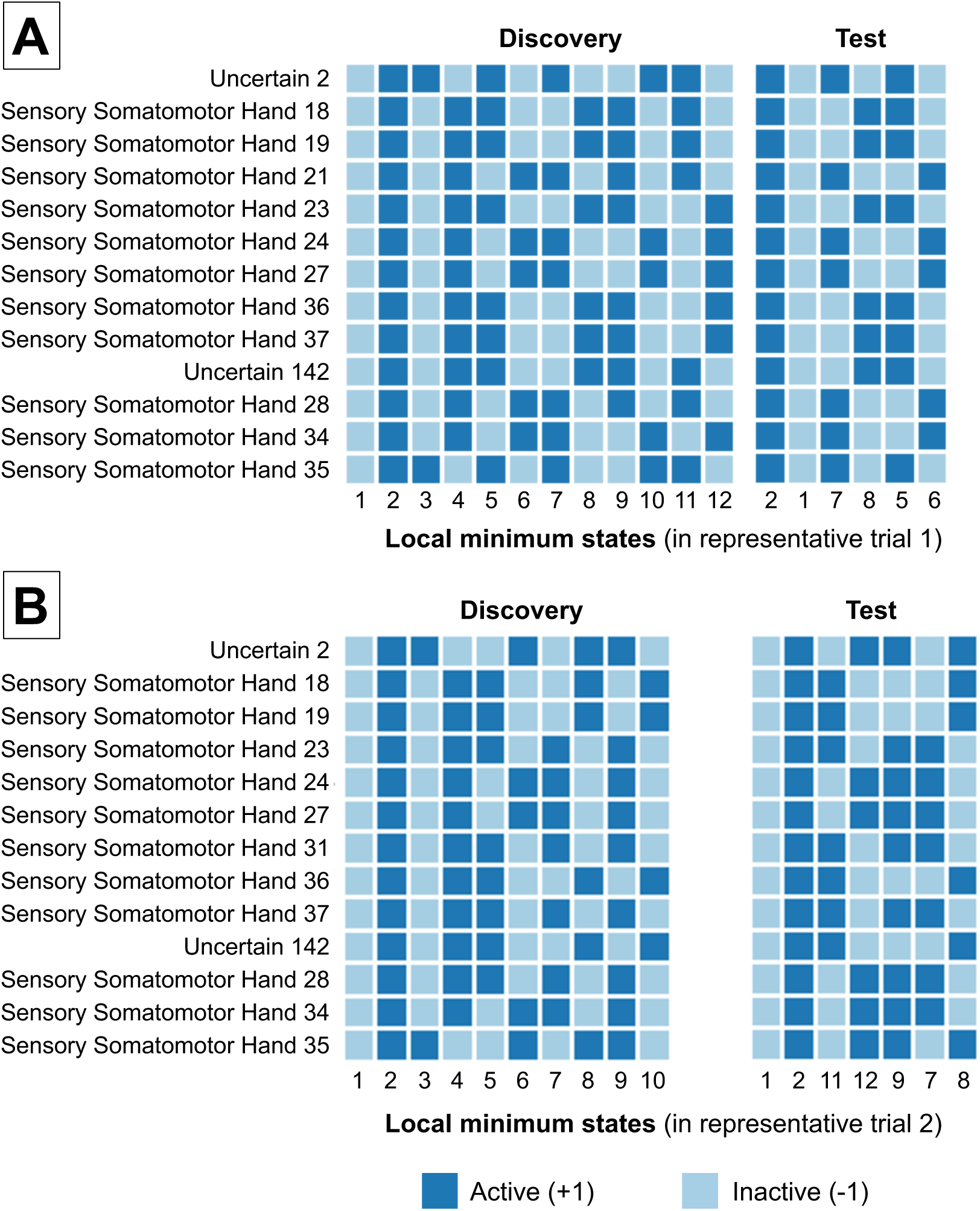
Observed local minimum states for representative trials: (A) trial 1 and (B) trial 2. . State numbers denote unique binary patterns; matching numbers indicate the same pattern across cohorts (if applicable).

**Table 4.** Frequency of local minimum occurrence in representative trial 1 (Significant results in bold; FDR-corrected *p*-value).

| State number | Cohort | probability of appearance |  | *p*-value (FDR corrected) | Effect size *r* |
| --- | --- | --- | --- | --- | --- |
|  |  | ASD | CTL |  |  |
| I | A | 0.367 | 0.361 | 0.436 | 0.0531 |
| 1 | B | 0.464 | 0.466 | **0.00386** | -0.276 |
| 2 | A | 0.349 | 0.344 | **0.00897** | 0.179 |
| 2 | B | 0.463 | 0.461 | **0.00386** | 0.285 |
| 3 | A | 0.106 | 0.111 | 0.0536 | -0.129 |
| 4 | A | 0.104 | 0.107 | 0.0888 | -0.114 |
| 5 | A | 0.0175 | 0.0176 | 0.432 | -0.0577 |
| 5 | B | 0.0156 | 0.0152 | 0.767 | -0.0266 |
| 6 | A | 0.0168 | 0.0174 | 0.498 | -0.0435 |
| 6 | B | 0.00662 | 0.00734 | **2.41E-10** | -0.588 |
| 7 | A | 0.00784 | 0.00846 | **0.00941** | -0.173 |
| 7 | B | 0.0297 | 0.0293 | 0.742 | -0.0446 |
| 8 | A | 0.00797 | 0.00820 | 0.0526 | -0.134 |
| 8 | B | 0.0210 | 0.0207 | 0.143 | 0.149 |
| 9 | A | 0.00665 | 0.00741 | **0.00897** | -0.194 |
| 10 | A | 0.00751 | 0.00801 | **0.00897** | -0.179 |
| 11 | A | 0.00562 | 0.00552 | 0.290 | 0.0760 |
| 12 | A | 0.00444 | 0.00460 | 0.699 | -0.0226 |

To validate the robustness of these findings, the same analysis was repeated on the independent validation dataset (Cohort B). In this cohort, six distinct local minimum states were observed Figure 10A, right panel), and notably, the local minimum state common to both cohorts (shared the same state number). These validation results supported the conclusion that the ASD group exhibited a higher probability of visiting a local minimum characterized by the simultaneous activation of all selected ROIs.

A second representative trial was similarly identified based on ROI selection consistency (Figure 10B). In this trial, 13 ROIs were selected, and 10 unique local minimum states were observed in Cohort A (Figure 10B, left panel). Mann-Whitney *U* tests revealed significant differences in the occurrence probability for seven states between groups (*p* < 0.05, FDR corrected; see Table 5). Again, several states were more frequently visited by the ASD group, while others were predominant in the CTL group.

**Table 5.** Frequency of local minimum occurrence in representative trial 2 (Significant results in bold; FDR-corrected p-values)

| State number | Cohort | probability of appearance |  | p-value (FDR corrected) | Effect size *r* |
| --- | --- | --- | --- | --- | --- |
|  |  | ASD | CTL |  |  |
| 1 | A | 0.369 | 0.365 | 0.540 | 0.0359 |
| 1 | B | 0.472 | 0.475 | **3.09E-05** | -0.396 |
| 2 | A | 0.355 | 0.351 | **0.00740** | 0.174 |
| 2 | B | 0.461 | 0.456 | **2.08E-08** | 0.529 |
| 3 | A | 0.0948 | 0.0984 | **0.0380** | -0.130 |
| 4 | A | 0.0950 | 0.0981 | 0.162 | -0.0851 |
| 5 | A | 0.0193 | 0.0202 | **0.0319** | -0.137 |
| 6 | A | 0.0197 | 0.0215 | **0.000559** | -0.235 |
| 7 | A | 0.0105 | 0.0109 | **0.007399** | -0.174 |
| 7 | B | 0.00863 | 0.00894 | **0.0167** | -0.240 |
| 8 | A | 0.0127 | 0.0127 | 0.0584 | 0.116 |
| 8 | B | 0.00828 | 0.00828 | 0.815 | -0.0386 |
| 9 | A | 0.0121 | 0.0109 | **0.00937** | 0.165 |
| 9 | B | 0.0137 | 0.0136 | 0.815 | 0.0347 |
| 10 | A | 0.0125 | 0.0117 | **0.00684** | 0.187 |
| 11 | B | 0.0203 | 0.0209 | 0.894 | -0.0121 |
| 12 | B | 0.0166 | 0.0174 | 0.815 | -0.0641 |

Validation on Cohort B confirmed the presence of seven local minima (Figure 10B, right panel), with overlapping state numbers across cohorts. Group differences in visiting specific states (notably States 2) were replicated, further supporting the generalizability of the observed ASD-specific dynamics.

### Scenarios 3 and 4

Supplementary Figures 8 and 10 show the convergence of the objective function across 1000 generations during 100 independents ELA/GAopt runs for the ASD and CTL groups, respectively. To evaluate the consistency of selected ROIs, binomial tests were performed on the selection results from each group’s 100 runs. Statistically significant ROIs after FDR correction are presented in Figure 11 and Supplementary Tables 4 (ASD) and 5 (CTL). The selection frequency distribution across all ROI candidates is shown in Supplementary Figures 9 (ASD) and 11 (CTL), reflecting how often each ROI was selected over 100 optimization trials. Similar to previous scenarios, ROI selection in Scenarios 3 and 4 showed high stability as shown in Figure 12 and Figure 13, respectively. In both the ASD and CTL groups, the pairwise Hamming distance between the 100 optimized sets (ROI selection vector) was significantly lower compared to the random set (Scenario 3: Mann- Whitney *U* test, *p* < 0.0001, FDR-corrected, Cohen’s *d* = 6.08, Figure 12A; Scenario 4: Mann-Whitney *U* test, *p* < 0.0001, FDR-corrected, Cohen’s *d* = 5.44, Figure 13A). The pairwise Jaccard index was significantly higher compared to the random set (Scenario 3: Mann-Whitney *U* test, *p* < 0.0001, FDR-corrected, Cohen’s *d* = 5.04, Figure 12B; Scenario 4: Mann-Whitney *U* test, *p* < 0.0001, FDR-corrected, Cohen’s *d* = 4.84, Figure 13B).

**Figure 11.**
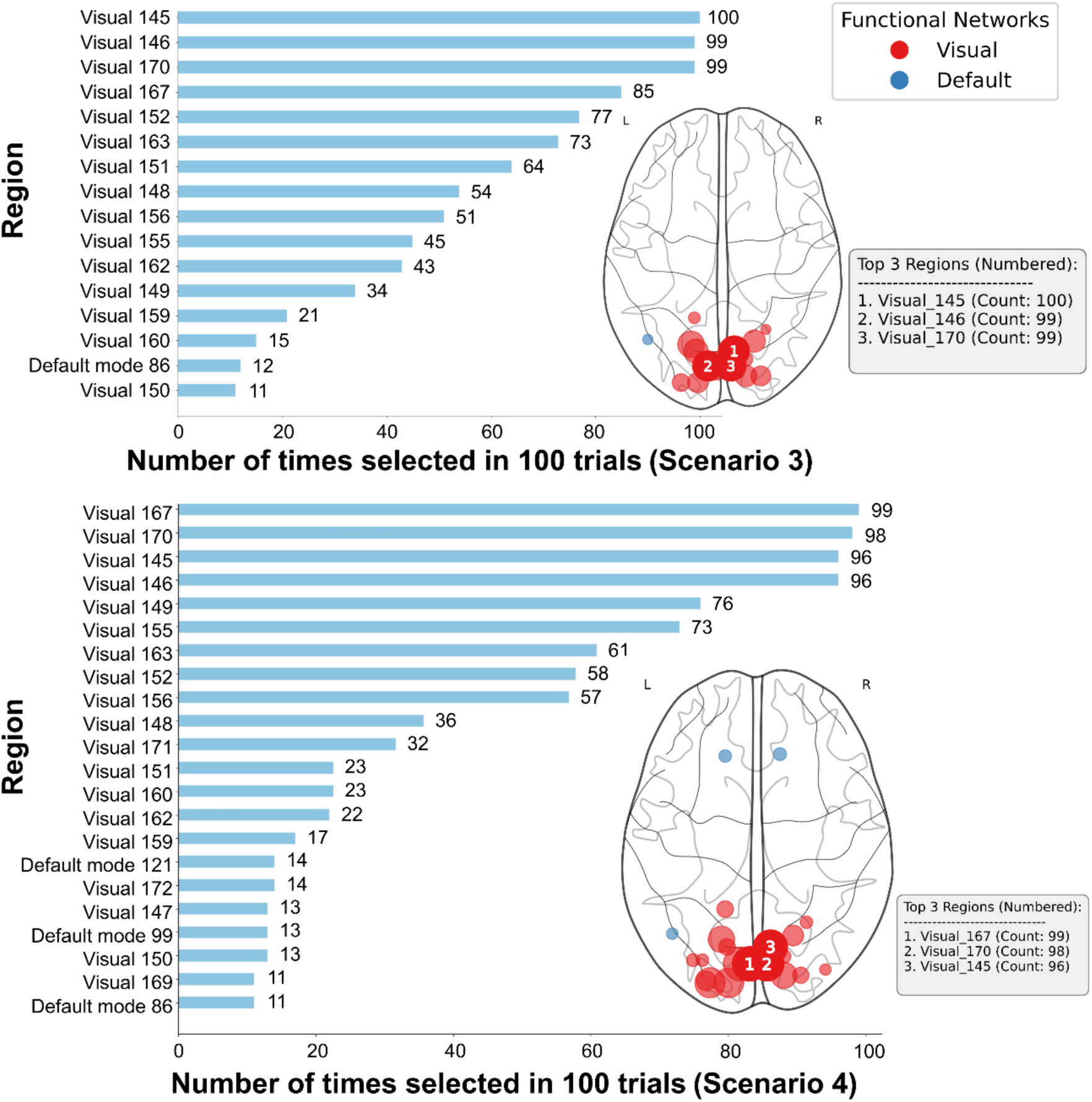
Selection frequency of each ROI across 100 runs of ELA/GAopt (Scenarios 3 and 4). The bar graph illustrates the frequency with which each ROI was selected across 100 independent optimization runs. A binomial test was performed to determine whether the frequency of each ROI exceeded chance levels. As a result, 16 ROIs were identified as statistically significant in Scenario 3 and 22 ROIs in Scenario 4 (*p* < 0.05, FDR-corrected). The ROIs are sorted in descending order of selection frequency. Their anatomical locations are also displayed on the glass brain plot. The circle size in the brain plot represents the frequency of each ROI’s selection, with larger circles indicating higher frequencies.

**Figure 12.**
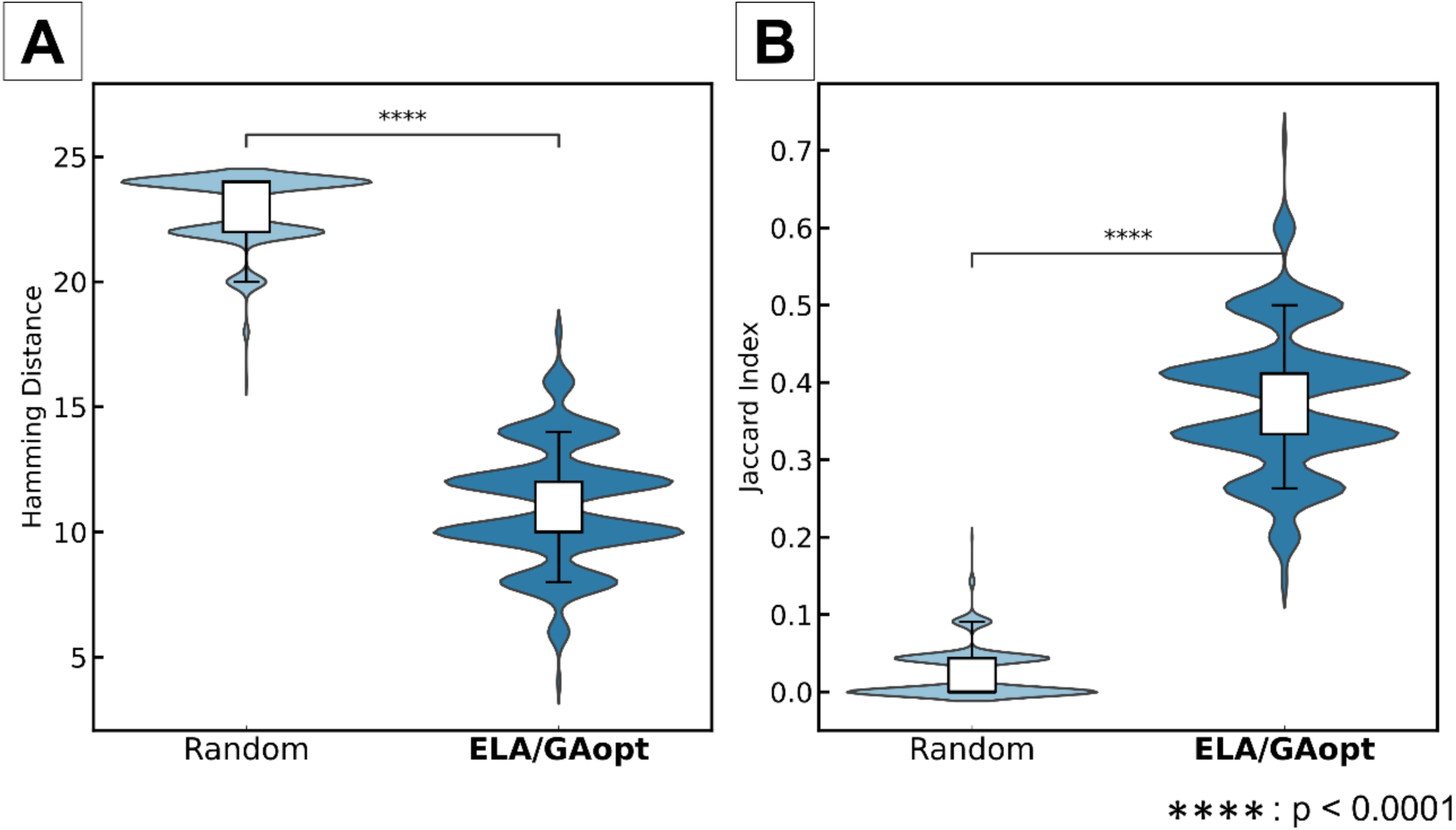
Stability evaluation of ROI selection using ELA/GAopt compared to random selection (Scenario 3). (**A**) The pairwise Hamming distances between ROI sets (ROI selection vectors). The pairwise Hamming distances between optimized ROI sets were significantly smaller than those between random sets (Mann-Whitney U test, *p* < 0.0001, FDR-corrected, Cohen’s *d* = 6.08). (**B**) The Jaccard index between ROI sets (ROI selection vectors). The pairwise Jaccard index between optimized ROI sets was significantly higher than that between randomly selected sets (Mann-Whitney U test, *p* < 0.0001, FDR-corrected, Cohen’s *d* = 5.04).

**Figure 13.**
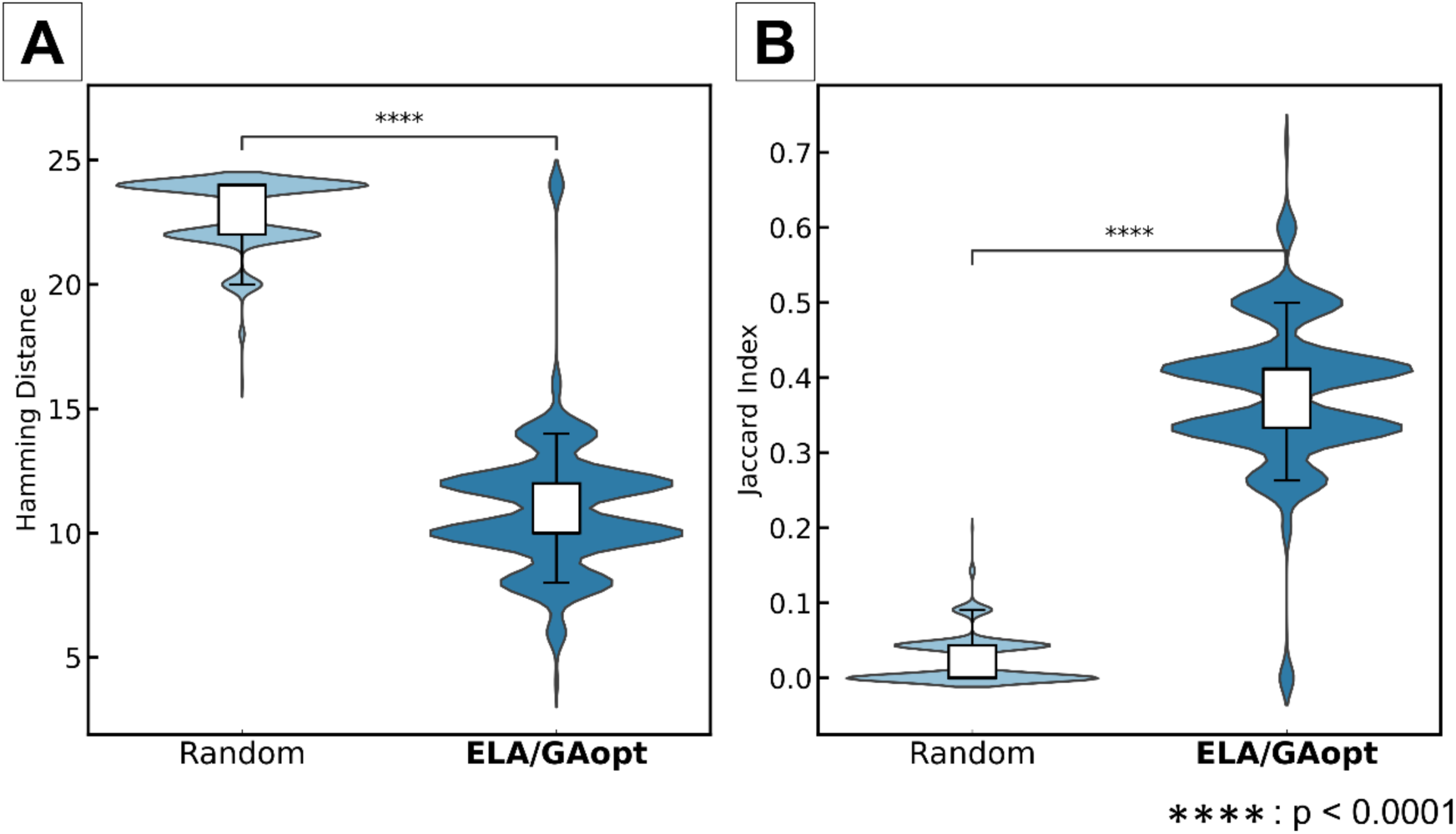
Stability evaluation of ROI selection using ELA/GAopt compared to random selection (Scenario 4). (**A**) The pairwise Hamming distances between ROI sets (ROI selection vectors). The pairwise Hamming distances between optimized ROI sets were significantly smaller than those between random sets (Mann-Whitney U test, *p* < 0.0001, FDR-corrected, Cohen’s *d* = 5.44). (**B**) The Jaccard index between ROI sets (ROI selection vectors). The pairwise Jaccard index between optimized ROI sets was significantly higher than that between randomly selected sets (Mann-Whitney U test, *p* < 0.0001, FDR-corrected, Cohen’s *d* = 4.84).

Using the ASD-optimized ROIs from Cohort C, we estimated energy landscapes for both ASD and CTL groups. The number of local minima obtained for each group was computed, and a group comparison was performed using the Mann-Whitney *U* test. The test revealed that the ASD group exhibited a significantly lower number of local minima than the CTL group (*U*_200_ = 4238.5, *p* = 0.043, effect size *r*: 0.13), suggesting that the ASD-optimized ROI configuration captures group-specific features of brain state dynamics.

Conversely, using the CTL-optimized ROIs from Cohort D, the CTL group exhibited significantly fewer local minima than the ASD group (*U*_200_ = 5924.5, *p* = 0.019, effect size *r*: 0.16), again suggesting that the CTL-optimized ROIs capture group-specific differences in brain state dynamics.

## Discussion

### Overview and advantages of the proposed method

The ELA/GAopt framework proposed in this study is designed as a versatile meta-framework that integrates energy landscape analysis with genetic algorithm-based feature selection. Its core architecture allows for the objective, data-driven identification of ROIs by optimizing a user-defined objective function tailored to specific research goals. In this study, we implemented a representative instance of this framework where the objective function was defined based on pMEM fitting accuracy and inter-individual variability of the personalized inverse temperature parameter. This specific configuration was chosen to enhance reproducibility and objectivity in brain dynamics modeling by ensuring model validity while capturing individual differences. However, it is important to emphasize that the ELA/GAopt framework itself is independent of these specific metrics; it can be flexibly extended to incorporate other criteria, such as correlations with clinical phenotypes or task performance, depending on the analysis requirements.

Traditional ROI selection approaches in neuroimaging studies often depend on established anatomical or functional priorities derived from previous research. While such hypothesis-driven selection improves consistency across studies, it may also introduce confirmation bias, limiting the analysis framework to existing knowledge and potentially missing emergent patterns not captured by prior findings.

In contrast, ELA/GAopt mitigates this bias by enabling fully data-driven ROI selection. This allows the model to explore brain region combinations not considered a priori, thereby expanding the search space for novel insights into brain dynamics. Such exploratory capacity is particularly crucial for advancing our understanding of complex and heterogeneous conditions, such as ASD. Hereafter, we discuss the results obtained in multiple scenarios to demonstrate the effectiveness and clinical applicability of our proposed ELA/GAopt.

In this context, it is important to clarify that the local minima identified by this framework correspond to metastable functional configurations, or attractor states, within the brain dynamics. These states represent specific patterns of recurrent co-activation among the selected ROIs that the system tends to revisit and maintain over time. Transitions between local minima reflect dynamic shifts in the functional organization of brain networks. Furthermore, it should be recognized that brain states derived from ELA are inherently constrained by the binarized pMEM framework. Thus, while local minima provide a powerful mathematical abstraction for characterizing macroscopic network dynamics, caution is warranted against over-interpreting these discrete states in strictly biological terms. Guided by this interpretive framework, we now assess the findings across multiple scenarios to clarify how ELA/GAopt supports the identification of both generalizable and condition-specific brain dynamics.

### Evaluation of reproducibility (Scenario 1)

Scenario 1 evaluated the generalizability of ELA/GAopt using the publicly available Creativity dataset (OpenNeuro: ds002330), which includes resting-state fMRI data. ELA/GAopt was applied to the discovery set, and the optimized ROI sets were subsequently applied to the test set. These ROI sets achieved similarly high objective function values in both datasets (Figure 4B, C, E, and F; *p* < 0.05, FDR-corrected), demonstrating that ELA/GAopt avoids overfitting and yields generalizable ROI selection.

Regarding the stochastic nature of GAs, we empirically validated the convergence stability through 100 independent runs using random initialization with different random seeds. The results demonstrated consistent convergence to high-fitness solutions with minimal variance across different seeds, supporting the robustness of the optimization process. Nevertheless, a more formal theoretical analysis of seed dependency remains a subject for future research to further characterize the algorithmic properties of ELA/GAopt.

It is also noteworthy that the effect sizes observed in the discovery set were remarkably large (Cohen’s *d* > 5). Unlike clinical effect sizes that measure biological differences between groups, the metric in this context quantifies the degree of separation achieved by optimization against a random baseline. Since the GA explicitly maximizes the objective function, it can produce objective values that are far from the distribution of random ROI selections, resulting in a large Cohen’s *d*. This should be interpreted as the magnitude of optimization gain relative to the random baseline, rather than a neurobiological effect.

Moreover, the similarity of local minimum states between the discovery and test sets was significantly higher (Hamming distance difference was smaller) for ROI sets selected by ELA/GAopt than for randomly selected ROIs (Figure 4A and D; *p* < 0.05, FDR-corrected). Because local minimum represents stable brain states, this result suggests ELA/GAopt effectively identifies ROI sets capturing stable, reproducible patterns.

Reproducibility is a critical issue in neuroimaging research, especially for clinical translation. Traditional ROI-based analyses may underestimate reproducibility because fMRI signals are inherently non-stationary and exhibit considerable inter-individual and temporal variability (Song et al., 2014, 2016). In contrast, ELA/GAopt potentially enhances repeatability by excluding noisy or irrelevant features during ROI selection. Importantly, the objective function of ELA/GAopt explicitly considers inter-individual variance (i.e., inverse temperature parameter of pMEM), which may further promote reproducibility in group-level energy landscape analyses. These improvements enhance the robustness of the findings and increase the potential for future clinical applications, including and personalized modeling of brain dynamics.

Our analysis using the high-dimensional HCP-YA dataset (*N* = 15, i.e., 2^15^ state space) revealed a significant trade-off between model fitting and stability of pattern reproducibility. ELA/GAopt achieved remarkably high reproducibility in objective function values with Cohen’s *d* > 6 in both discovery and test sets, confirming the algorithm’s robustness in maximizing pMEM fit and variance of the personalized inverse temperature parameter. However, the structural reproducibility of local minima states, specifically the Hamming distance between local minima, did not exhibit a significant advantage over random selection.

Detailed post-hoc validation indicated that optimized ROI sets consistently identified a significantly higher number of local minima compared to random sets. This suggests that ELA/GAopt captures more complex energy landscapes reflecting subtle dynamic transitions, whereas random selection tends to yield flatter, simpler landscapes with fewer states. In high-dimensional spaces, this increased complexity inherently makes the exact replication of specific local minimum patterns more challenging (i.e., the “curse of dimensionality”). Consequently, while ELA/GAopt ensures highly optimized model fitting, researchers should recognize that the precise configuration of local minima may exhibit greater cross-dataset variability in high-dimensional settings (*N* ≥ 15) compared to lower-dimensional configurations (*N* = 10). This highlights a fundamental trade-off between maximizing model accuracy and maintaining the structural stability of the energy landscape.

Furthermore, the ROIs selected by ELA/GAopt were primarily centered in the occipital and somatomotor networks, as well as core components of the default mode network (DMN) such as the angular gyrus, precuneus, and inferior temporal lobe. We acknowledge that the selection of these primary sensory-motor and visual regions is, to some extent, an expected consequence of the mathematical characteristics of pMEM. Since participants in resting-state fMRI often engage in subtle visual processing or motor activity (e.g., small hand movements), these regions naturally exhibit high neural variability (Koba et al., 2021; Mao et al., 2015), which can contribute to higher model fitting accuracy. Therefore, the consistent identification of these areas serves as a basic validation that ELA/GAopt effectively captures the major sources of variance within the data. This is consistent with recent evidence suggesting that motor-related regions exhibit significant coordinated activity even during the resting state (Seitzman et al., 2019; J. A. Smith et al., 2023).

It is important to emphasize that the objective function employed in this study serves as a representative implementation to demonstrate the framework’s fundamental utility. As ELA/GAopt is designed as a versatile meta-framework independent of specific metrics, it can be flexibly extended to incorporate diverse optimization criteria, depending on the research objective. Crucially, however, even within the current implementation, ELA/GAopt is not a mere ‘variance maximizer.’ If maximizing fitting accuracy alone were the goal, the algorithm might converge toward selecting trivial high-amplitude regions or noise artifacts. To prevent such a result, our objective function specifically incorporates a term representing the inter-individual variability of the personalized inverse temperature parameter (*β*). This design ensures that the identified ROI sets, including the DMN regions associated with self-referential thought and memory integration, are selected not only for their fitting accuracy to the group-level model but also for their ability to capture individual differences in brain dynamics. By balancing model fidelity with individual variability, ELA/GAopt enables the data-driven discovery of ROI combinations that reflect complex collective dynamics rather than simple recording-condition manifestations.

### Detection of ASD-specific dynamics (Scenarios 2–4)

Scenarios 2, 3, and 4 employed the ABIDE II dataset to investigate whether ELA/GAopt could detect ASD-specific features in brain state dynamics.

In Scenario 2, ROIs selected by ELA/GAopt were mainly located within the sensory somatomotor hand network (Figure 8). Significant group differences in specific local minima probability were observed between the ASD and typically developing (CTL) groups (Table 4, Table 5 and Figure 10; *p* < 0.05, FDR-corrected). Notably, ASD participants more frequently visited local minimum characterized by the co-activation of all selected ROIs. This dynamic signature was replicated in an independent validation cohort (Cohort B), providing support for its robustness.

These findings are consistent with prior reports that identified local overconnectivity and long-range underconnectivity as characteristic features of brain organization in individuals with ASD (Fu et al., 2025). The predominance of globally ‘active’ local minimum (e.g., State 2 identified in Scenario 2, where all selected ROIs are in the active state +1) suggests excessive synchronization of neural activity in ASD, potentially reducing the flexibility of brain state transitions. This observation aligns with previous ELA studies showing overstability of neural dynamics (Watanabe & Rees, 2017), and increased functional connectivity among visual and sensorimotor networks in infants with ASD, which correlated with symptom severity (Chen et al., 2021). Moreover, manifold learning studies identified connectivity patterns in sensorimotor networks as discriminative features for distinguishing ASD from schizophrenia (Hao et al., 2024).

Taken together, these results suggest that ASD is associated with a distinct neurodynamic profile characterized by reduced state variability and increased co-activation within specific functional networks. The ELA/GAopt demonstrated its effectiveness in uncovering such network-specific dynamic biomarkers in a fully data-driven manner.

Scenario 3 further evaluated this hypothesis by applying the ROI set optimized for the ASD group to the CTL group. The ROI sets optimized for the ASD group primarily consisted of visual network regions, as indicated in Figure 11. The application of the ASD-optimized ROIs to the ELA on the CTL group led to a significant increase in the number of observed local minima in CTL participants (*p* < 0.05), indicating that ASD-optimized ROIs poorly capture the neurodynamic patterns of typically developing individuals. This supports previous work suggesting that ASD is characterized by atypical over-coupling of visual networks, which may limit brain state transitions (Sadeghian et al., 2021; Yamasaki et al., 2017).

Scenario 4 provided a complementary test by applying CTL-optimized ROIs to the ASD group. Similarly, the number of local minima observed in ASD participants was significantly increased (*p* < 0.05). Considering the results in Scenario 3, they suggest that the two groups exhibit fundamentally different brain state dynamics and network configurations. This further strengthens previous findings that ASD and CTL groups possess distinct neurodynamic architectures (Chen et al., 2021; Fu et al., 2025; Hao et al., 2024; Kozhemiako et al., 2020; Watanabe & Rees, 2017; Yamasaki et al., 2017).

These inter-application results suggest that ELA/GAopt can identify ROI sets that are not only discriminative but also specific to particular neurodevelopmental conditions. This highlights its potential utility for characterizing condition-specific brain state dynamics, serving as a steppingstone for future biomarker research. While these findings warrant further validation across diverse and larger independent clinical cohorts to ensure clinical utility, our proposed ELA/GAopt demonstrates strong potential as a data-driven framework for identifying meaningful brain dynamics and holds promise for future applications in both basic and translational neuroscience research.

### Comparison with other ROI selection strategies

It is essential to distinguish ELA/GAopt from other data-driven dimensionality reduction methods commonly employed in neuroimaging, such as ICA-based parcellation, PCA (Calhoun et al., 2009; Guo & Pagnoni, 2008; Seewoo et al., 2021; Varoquaux et al., 2010) and clustering-based approaches (Craddock et al., 2011; Thirion et al., 2014). Although these techniques effectively reduce data complexity, they typically generate spatial maps or components that do not align with established anatomical boundaries, complicating the biological interpretation of the resulting energy landscape nodes. In contrast, ELA/GAopt performs subset selection within predefined anatomical atlas spaces, such as the Dosenbach or Power atlases used in the current study. This approach ensures that the selected nodes retain their original anatomical labels (e.g., “posterior cingulate cortex”), thereby maintaining high interpretability and facilitating direct comparisons with existing literature. While this study demonstrates the utility of the framework, rigorous comparative work evaluating energy landscapes derived from GA-selected subsets versus ICA- or clustering-based components remains an important next step for validation.

Furthermore, ELA should be distinguished from other dynamic modeling approaches, such as dynamic functional connectivity (dFC) or dynamic graph theory. Whereas dFC often relies on sliding-window correlations to track continuous fluctuations in coupling strength (Hindriks et al., 2016), ELA models brain dynamics as transitions between discrete, energy-defined metastable states within a global landscape. ELA/GAopt contributes to this paradigm by providing a data-driven method to identify the specific ROI combinations that are well-suited to this energy-based representation under a user-defined objective. Moreover, the inherent flexibility of our meta-framework allows future studies to integrate graph-theoretic metrics, such as global efficiency or modularity, directly into the objective function. This could help bridge the gap between energy-based and connectivity-based perspectives on macroscopic brain dynamics.

Finally, unlike standard feature selection techniques such as recursive feature elimination (RFE), which primarily aim to maximize predictive accuracy for external labels (Bulut et al., 2025; Vafaie & Imam, n.d.), ELA/GAopt is a model-dependent selection framework. Its primary objective is to satisfy the specific mathematical constraints of the pairwise maximum entropy model (pMEM), notably the limitation of 10-15 ROIs, while maximizing (or minimizing) the objective function specified by researchers based on their research objective. Given the non-linear nature of pMEM fitting, in which complex interactions between nodes determine system energy, heuristic global optimization via genetic algorithms is theoretically superior to greedy methods that may overlook critical interactions. Although direct performance comparisons with other reduction strategies are left for future work, this study establishes the fundamental utility of GA-based optimization for biologically interpretable energy landscape analysis.

### Methodological limitations and future directions

While this study introduced ELA/GAopt as a novel data-driven method for ROI selection and demonstrated its utility across multiple neuroimaging datasets, several methodological limitations should be acknowledged.

First, although genetic algorithms (GAs) are well suited for nonlinear and high-dimensional optimization problems, they are computationally more expensive than commonly used gradient-based algorithms. While GAs has the advantage of conducting global search through multipoint exploration, they require numerous objective function evaluations—in this case, involving repeated pMEM fitting steps. For example, 10,000,000 function evaluations were required across 100 trials of ELA/GAopt in Scenario 1, resulting in a total computation time of 100,000 seconds. Because the necessary computational effort depends on the problem’s complexity, analysts must determine suitable termination criteria (e.g., the number of generations) empirically, which can be burdensome. In this study, we adopted conservative parameter setting (1,000 generations across 100 independent runs) to rigorously validate the convergence and stability of the algorithm. However, for practical applications, such extensive iterations may be reduced once solution stability has been empirically established. Furthermore, genetic algorithms are inherently parallelizable: because the evaluation of each individual within a population (i.e., pMEM fitting for a candidate ROI set) is computationally independent, computation scales approximately linearly with the number of available CPU or GPU cores. Future implementations that leverage high-performance computing clusters could substantially reduce execution time, thereby facilitating the application of ELA/GAopt to even larger datasets.

Second, the objective function used in this study was defined as the weighted sum of two components: pMEM fitting accuracy and the inter-individual variance of the inverse temperature parameter. While this single-objective formulation, with a weighted sum of multiobjective components, provides a practical means to combine multiple objectives, it requires the analyst to determine the relative importance of each component a priori. Moreover, controlling the importance is often not easy. A more flexible approach would be to adopt a multiobjective optimization framework that simultaneously optimizes each objective without requiring predefined weights. This strategy would explicitly characterize the trade-offs between competing goals and generate a Pareto front of optimal solutions, from which the most appropriate configuration could be selected based on study-specific needs. Incorporating multiobjective optimization into ELA/GAopt in future work may improve the robustness and diversity of the automated ROI selection.

Third, for the multi-site ABIDE II dataset, explicit data harmonization techniques such as ComBat (J.-P. Fortin et al., 2018) were not applied directly to the BOLD time-series data. Site-related variability is a well-recognized confounder in multisite neuroimaging studies. However, standard harmonization methods are typically optimized for summary statistics, such as functional connectivity matrices or cortical thickness, rather than raw time-series data (J.-P. Fortin et al., 2017; Y.-W. Wang et al., 2023). Directly applying such corrections to BOLD signals carries the risk of distorting temporal autocorrelation structures and nonlinear dynamics that are essential for ELA (Sahoo & Davatzikos, 2021; Saponaro et al., 2022). Instead, we mitigated the risk of site-specific overfitting through a rigorous site-disjoint validation design. As shown in Table 3, the discovery set (Cohort A) and validation set (Cohort B) consisted of completely non-overlapping sites (e.g., NYU and UCLA in Cohort A; OHSU and USM in Cohort B). The replication of the identified ROI sets and brain dynamics in Cohort B suggests that our results are less likely to be driven solely by site-specific artifacts; nevertheless, residual site effects cannot be ruled out without explicit harmonization. Although standard denoising was performed, future work should investigate advanced harmonization frameworks specifically designed for dynamic time-series analysis to further reduce inter-site variability.

Furthermore, this study accounted for inter-individual variation by incorporating the variance of the inverse temperature parameter (*β*) into the objective function. We acknowledge that attributing inter-individual differences solely to *β* is a strong assumption. As suggested by recent literature, individual variability is also likely driven by structural differences in the energy landscape itself, such as the number and configuration of local minima. In the current framework, the *β*-variance term was employed primarily as a computational heuristic to prevent the algorithm from converging toward trivial solutions that maximize fitting accuracy while ignoring subject-level diversity. Future research should explore alternative objective functions that directly quantify topological differences in energy landscapes across individuals.

Finally, while the ASD-related findings in Scenarios 2-4 are promising, they should be interpreted with caution. The identification of group-specific dynamics in the ABIDE II dataset serves primarily as a representative application of the ELA/GAopt framework, rather than as evidence for definitive clinical biomarkers suitable for immediate diagnostic use. The primary value of our method lies in its ability to objectively identify candidate ROI subsets and landscape-derived features (e.g., state transition probabilities) that characterize condition-specific dynamics, thereby providing a reproducible pathway for feature selection in future diagnostic modeling. Although our findings were validated using an independent cohort within the same dataset, further replication in completely independent, larger-scale cohorts is necessary to ensure generalizability to the broader ASD population. Future studies should also investigate how these dynamic signatures relate to specific clinical symptoms or behavioral measures.

Despite these methodological limitations, our results demonstrate that ELA/GAopt offers a powerful and generalizable approach for exploring brain dynamics in a fully data-driven manner, with potential for broader applications in neuroscience research.

## Data Availability

All the datasets used in this study are publicly available. The source codes of the proposed method are available on our GitHub repository (https://github.com/MIS-Lab-Doshisha/ela-gaopt).

## Declaration of generative AI and AI-assisted technologies in the writing process

During the preparation of this work the authors used ChatGPT and NotebookLM in order to improve the readability and language of the manuscript (e.g., clarity, conciseness, and writing style). After using this tool/service, the authors reviewed and edited the content as needed and take full responsibility for the content of the published article.

## Acknowledgements

This work was supported by JSPS KAKENHI Grant Number JP24K03028, and in part by grants-in-aid from the Harris Science Research Institute of Doshisha University. We acknowledge the significant contributions of ABIDE-II and its primary source of funding: NIMH 5R21MH107045. HCP-YA data were provided by the Human Connectome Project, WU-Minn Consortium (Principal Investigators: David Van Essen and Kamil Ugurbil; 1U54MH091657) funded by the 16 NIH Institutes and Centers that support the NIH Blueprint for Neuroscience Research; and by the McDonnell Center for Systems Neuroscience at Washington University.

