## Supplementary for "Energy Landscape Analysis with Automated Region-of-Interest Selection via Genetic Algorithms"

### Supplementary Materials

#### Supplementary Table 1. Brain regions significantly selected in ELA/GAopt (Scenario 1, on Creativity data; Significant results in bold; FDR-corrected p-values)

| | | | | Data validation | | | |
|---|---|---|---|---|---|---|---|
| Region | Count | FDR-corrected p | Cohen's h | x | y | z | Network |
| occipital 142 | 99 | **<1E-10** | 2.436 | -29 | -75 | 28 | occipital |
| angular gyrus 117 | 92 | **<1E-10** | 2.063 | 51 | -59 | 34 | default |
| post occipital 153 | 88 | **<1E-10** | 1.929 | 29 | -81 | 14 | occipital |
| occipital 139 | 83 | **<1E-10** | 1.786 | 29 | -73 | 29 | occipital |
| dACC 27 | 40 | **<1E-10** | 0.864 | 9 | 20 | 34 | cingulo-opercular |
| post occipital 156 | 38 | **<1E-10** | 0.823 | -37 | -83 | -2 | occipital |
| mid insula 44 | 36 | **<1E-10** | 0.781 | 37 | -2 | -3 | cingulo-opercular |
| aPFC 3 | 35 | **<1E-10** | 0.761 | -29 | 57 | 10 | fronto-parietal |
| vFC 40 | 31 | **<1E-10** | 0.676 | -48 | 6 | 1 | cingulo-opercular |
| aPFC 8 | 29 | **<1E-10** | 0.632 | 27 | 49 | 26 | cingulo-opercular |
| precentral gyrus 67 | 28 | **<1E-10** | 0.610 | -54 | -22 | 22 | sensorimotor |
| precuneus 105 | 28 | **<1E-10** | 0.610 | 5 | -50 | 33 | default |
| frontal 32 | 24 | **5.40E-08** | 0.519 | 58 | 11 | 14 | sensorimotor |
| inf temporal 91 | 24 | **5.40E-08** | 0.519 | -61 | -41 | -2 | default |
| angular gyrus 102 | 21 | **4.55E-06** | 0.447 | -41 | -47 | 29 | cingulo-opercular |
| temporal 82 | 18 | **0.000230** | 0.371 | -41 | -37 | 16 | sensorimotor |
| med cerebellum 120 | 17 | **0.000723** | 0.345 | -6 | -60 | -15 | cerebellum |
| post occipital 154 | 13 | **0.0428** | 0.232 | 33 | -81 | -2 | occipital |

**Supplementary Table 2. Brain regions significantly selected in ELA/GAopt (Scenario 1, on HCP-YA data; Significant results in bold; FDR-corrected p-values)**

| | | | | Data validation | | | |
|---|---|---|---|---|---|---|---|
| Region | Count | FDR-corrected p | Cohen's h | x | y | z | Network |
| Visual_155 | 96 | **<1E-10** | 2.258 | -14 | -91 | 31 | Visual |
| Visual_162 | 92 | **<1E-10** | 2.087 | 24 | -87 | 24 | Visual |
| Visual_145 | 86 | **<1E-10** | 1.893 | 8 | -72 | 11 | Visual |
| Visual_149 | 85 | **<1E-10** | 1.864 | -24 | -91 | 19 | Visual |
| Sensory_Somatomotor_Mouth_44 | 83 | **<1E-10** | 1.810 | 51 | -6 | 32 | Sensory |
| Sensory_Somatomotor_Mouth_42 | 82 | **<1E-10** | 1.784 | -49 | -11 | 35 | Sensory |
| Visual_170 | 80 | **<1E-10** | 1.733 | 6 | -81 | 6 | Visual |
| Visual_146 | 75 | **<1E-10** | 1.613 | -8 | -81 | 7 | Visual |
| Sensory_Somatomotor_Mouth_45 | 73 | **<1E-10** | 1.567 | -53 | -10 | 24 | Sensory |
| Visual_167 | 72 | **<1E-10** | 1.545 | -3 | -81 | 21 | Visual |
| Visual_156 | 62 | **<1E-10** | 1.332 | 15 | -87 | 37 | Visual |
| Sensory_Somatomotor_Hand_27 | 51 | **<1E-10** | 1.109 | -38 | -27 | 69 | Sensory |
| Sensory_Somatomotor_Hand_24 | 43 | **<1E-10** | 0.949 | -40 | -19 | 54 | Sensory |
| Sensory_Somatomotor_Hand_23 | 40 | **<1E-10** | 0.888 | -23 | -30 | 72 | Sensory |
| Visual_152 | 37 | **<1E-10** | 0.826 | -18 | -68 | 5 | Visual |
| Sensory_Somatomotor_Hand_46 | 26 | **4.62E-10** | 0.589 | 66 | -8 | 25 | Sensory |
| Visual_159 | 23 | **7.31E-08** | 0.519 | 15 | -77 | 31 | Visual |
| Sensory_Somatomotor_Hand_34 | 21 | **1.61E-06** | 0.471 | -21 | -31 | 61 | Sensory |
| Visual_163 | 16 | **0.0014** | 0.342 | 6 | -72 | 24 | Visual |
| Visual_157 | 15 | **0.0043** | 0.314 | 29 | -77 | 25 | Visual |

**Supplementary Table 3. Brain regions significantly selected in ELA/GAopt (Scenario 2; Significant results in bold; FDR-corrected p-values).**

| | | | | Data validation | | | |
|---|---|---|---|---|---|---|---|
| Region | Count | FDR-corrected p | Cohen's h | x | y | z | Network |
| Sensory_Somatomotor_Hand_24 | 98 | **<1E-10** | 2.410 | -40 | -19 | 54 | Sensory |
| Sensory_Somatomotor_Hand_27 | 98 | **<1E-10** | 2.410 | -38 | -27 | 69 | Sensory |
| Sensory_Somatomotor_Hand_37 | 97 | **<1E-10** | 2,346 | -38 | -15 | 69 | Sensory |
| Uncertain_142 | 93 | **<1E-10** | 2.159 | -12 | -95 | 13 | Uncertain |
| Uncertain_2 | 92 | **<1E-10** | 2.121 | 27 | -97 | -13 | Uncertain |
| Sensory_Somatomotor_Hand_19 | 88 | **<1E-10** | 1.987 | 13 | -33 | 75 | Sensory |
| Sensory_Somatomotor_Hand_18 | 85 | **<1E-10** | 1.899 | -7 | -33 | 72 | Sensory |
| Sensory_Somatomotor_Hand_34 | 82 | **<1E-10** | 1.818 | -21 | -31 | 61 | Sensory |
| Sensory_Somatomotor_Hand_28 | 78 | **<1E-10** | 1.718 | 20 | -29 | 60 | Sensory |
| Sensory_Somatomotor_Hand_21 | 68 | **<1E-10** | 1.492 | 29 | -17 | 71 | Sensory |
| Sensory_Somatomotor_Hand_36 | 66 | **<1E-10** | 1.449 | 42 | -20 | 55 | Sensory |
| Sensory_Somatomotor_Hand_23 | 62 | **<1E-10** | 1.366 | -23 | -30 | 72 | Sensory |
| Sensory_Somatomotor_Hand_31 | 49 | **<1E-10** | 1.103 | 10 | -17 | 74 | Sensory |
| Sensory_Somatomotor_Hand_35 | 31 | **<1E-10** | 0.733 | -13 | -17 | 75 | Sensory |

**Supplementary Table 4. Brain regions significantly selected in ELA/GAopt (Scenario 3; Significant results in bold; FDR-corrected p-values).**

| | | | | Data validation | | | |
|---|---|---|---|---|---|---|---|
| Region | Count | FDR-corrected p | Cohen's h | x | y | z | Network |
| Visual_145 | 100 | **<1E-10** | 2.712 | 8 | -72 | 11 | Visual |
| Visual_146 | 99 | **<1E-10** | 2.512 | -8 | -81 | 7 | Visual |
| Visual_170 | 99 | **<1E-10** | 2.512 | 6 | -81 | 21 | Visual |
| Visual_167 | 85 | **<1E-10** | 1.916 | -3 | -81 | 21 | Visual |
| Visual_152 | 77 | **<1E-10** | 1.712 | -18 | -68 | 5 | Visual |
| Visual_163 | 73 | **<1E-10** | 1.619 | 6 | -72 | 24 | Visual |
| Visual_151 | 64 | **<1E-10** | 1.425 | -15 | -72 | -8 | Visual |
| Visual_148 | 54 | **<1E-10** | 1.221 | 20 | -66 | 2 | Visual |
| Visual_156 | 51 | **<1E-10** | 1.161 | 15 | -87 | 37 | Visual |
| Visual_155 | 45 | **<1E-10** | 1.041 | -14 | -91 | 31 | Visual |
| Visual_162 | 43 | **<1E-10** | 1.001 | 24 | -87 | 24 | Visual |
| Visual_149 | 34 | **<1E-10** | 0.815 | -24 | -91 | 19 | Visual |
| Visual_159 | 21 | **3.92E-08** | 0.522 | 15 | -77 | 31 | Visual |
| Visual_160 | 15 | **0.000428** | 0.366 | -16 | -52 | -1 | Visual |
| DefaultMode_86 | 12 | **0.0168** | 0.278 | -44 | -65 | 35 | Default |
| Visual_150 | 11 | **0.0463** | 0.246 | 27 | -59 | -9 | Visual |

**Supplementary Table 5. Brain regions significantly selected in ELA/GAopt (Scenario 4; Significant results in bold; FDR-corrected p-values).**

| | | | | Data validation | | | |
|---|---|---|---|---|---|---|---|
| Region | Count | FDR-corrected p | Cohen's h | x | y | z | Network |
| Visual_167 | 99 | **<1E-10** | 2.512 | -3 | -81 | 21 | Visual |
| Visual_170 | 98 | **<1E-10** | 2.428 | 6 | -81 | 6 | Visual |
| Visual_145 | 96 | **<1E-10** | 2.309 | 8 | -72 | 11 | Visual |
| Visual_146 | 96 | **<1E-10** | 2.309 | -8 | -81 | 7 | Visual |
| Visual_149 | 76 | **<1E-10** | 1.688 | -24 | -91 | 19 | Visual |
| Visual_155 | 73 | **<1E-10** | 1.619 | -14 | -91 | 31 | Visual |
| Visual_163 | 61 | **<1E-10** | 1.363 | 6 | -72 | 24 | Visual |
| Visual_152 | 58 | **<1E-10** | 1.302 | -18 | -68 | 5 | Visual |
| Visual_156 | 57 | **<1E-10** | 1.282 | 15 | -87 | 37 | Visual |
| Visual_148 | 36 | **<1E-10** | 0.857 | 20 | -66 | 2 | Visual |
| Visual_171 | 32 | **<1E-10** | 0.773 | -26 | -90 | 3 | Visual |
| Visual_151 | 23 | **1.13E-09** | 0.571 | -15 | -72 | -8 | Visual |
| Visual_160 | 23 | **1.13E-09** | 0.571 | -16 | -52 | -1 | Visual |
| Visual_162 | 22 | **6.55E-09** | 0.547 | 24 | -87 | 24 | Visual |
| Visual_159 | 17 | **2.42E-05** | 0.420 | 15 | -77 | 31 | Visual |
| DefaultMode_121 | 14 | **0.00141** | 0.337 | 13 | 30 | 59 | Default |
| Visual_172 | 14 | **0.00141** | 0.337 | -33 | -79 | -13 | Visual |
| Visual_147 | 13 | **0.00416** | 0.308 | -28 | -79 | 19 | Visual |
| DefaultMode_99 | 13 | **0.00416** | 0.308 | -16 | 29 | 53 | Default |
| Visual_150 | 13 | **0.00416** | 0.308 | 27 | -59 | -9 | Visual |
| Visual_169 | 11 | **0.0358** | 0.246 | 37 | -84 | 13 | Visual |
| DefaultMode_86 | 11 | **0.0358** | 0.246 | -44 | -65 | 35 | Default |

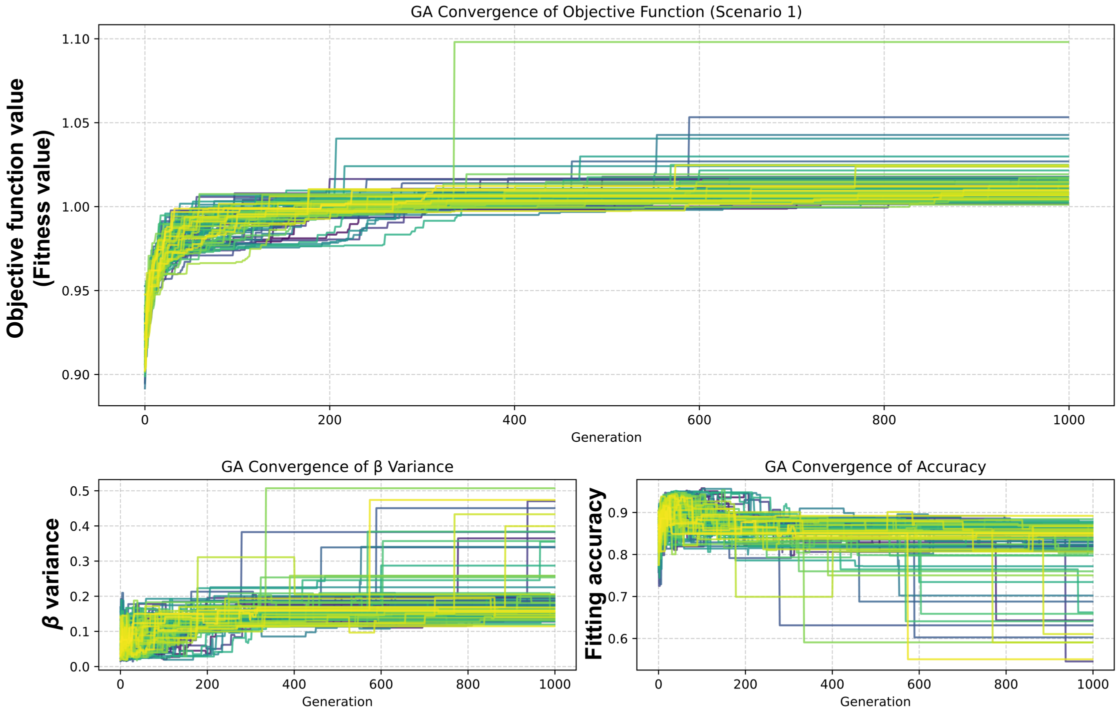


**Supplementary Figure 1. Convergence of the objective function with ELA/GAopt over 1000 generations for 100 runs with different starting population (Scenario 1, on Creativity data).** The objective function (top figure) is the sum of the β variance (bottom left) and the pMEM fitting accuracy (bottom right).

ELA with Genetic ROI Optimization

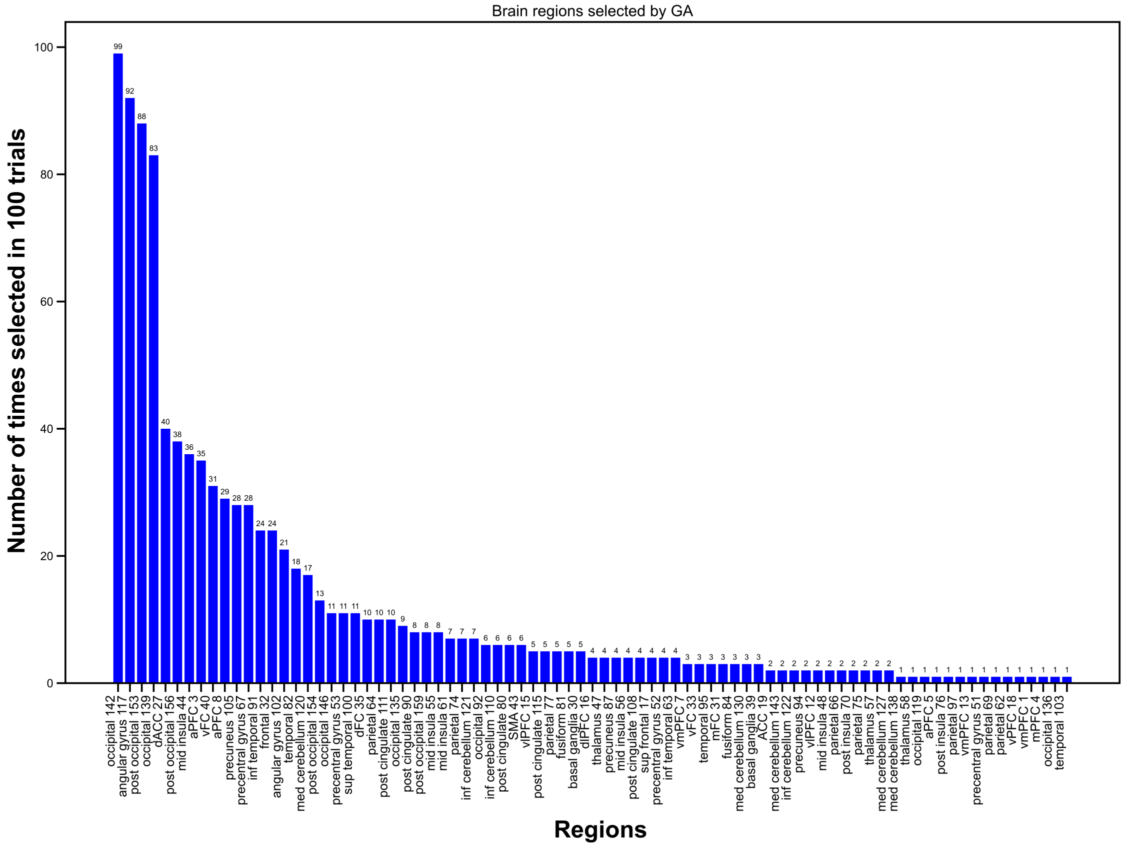


**Supplementary Figure 2. Selection frequency of each ROI across 100 runs of ELA/GAopt (Scenario 1, Creativity data).**

A

| Number | ROI label | Number | ROI label |
|---|---|---|---|
| 0 | med cerebellum 120 | 5 | occipital 142 |
| 1 | aPFC 8 | 6 | occipital 139 |
| 2 | vFC 40 | 7 | post occipital 156 |
| 3 | angular gyrus 117 | 8 | post occipital 153 |
| 4 | inf temporal 91 | 9 | frontal 32 |

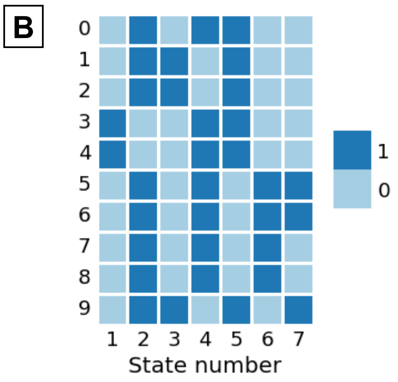


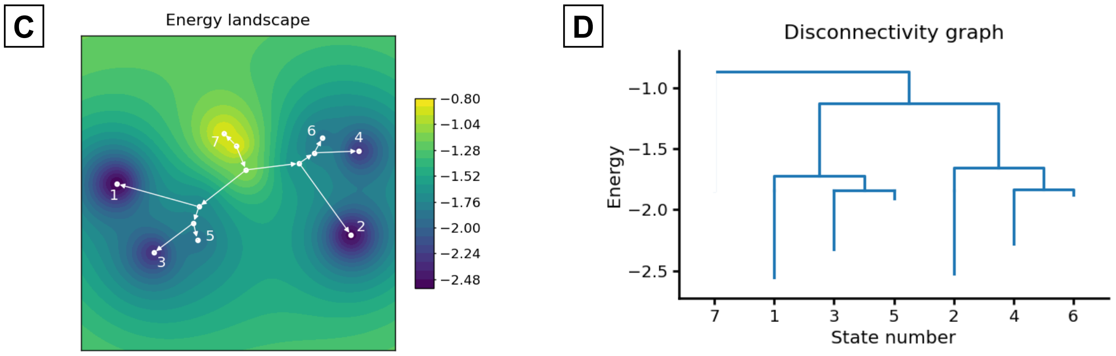


**Supplementary Figure 3. Example of energy landscape analysis using the optimized ROI set specified by a representative run consisting solely of the 18 significantly selected ROIs.** (A) List of the selected ROIs. (B) Observed local minimum states. (C) The resulting energy landscape. (D) Disconnectivity graph of the energy landscape.

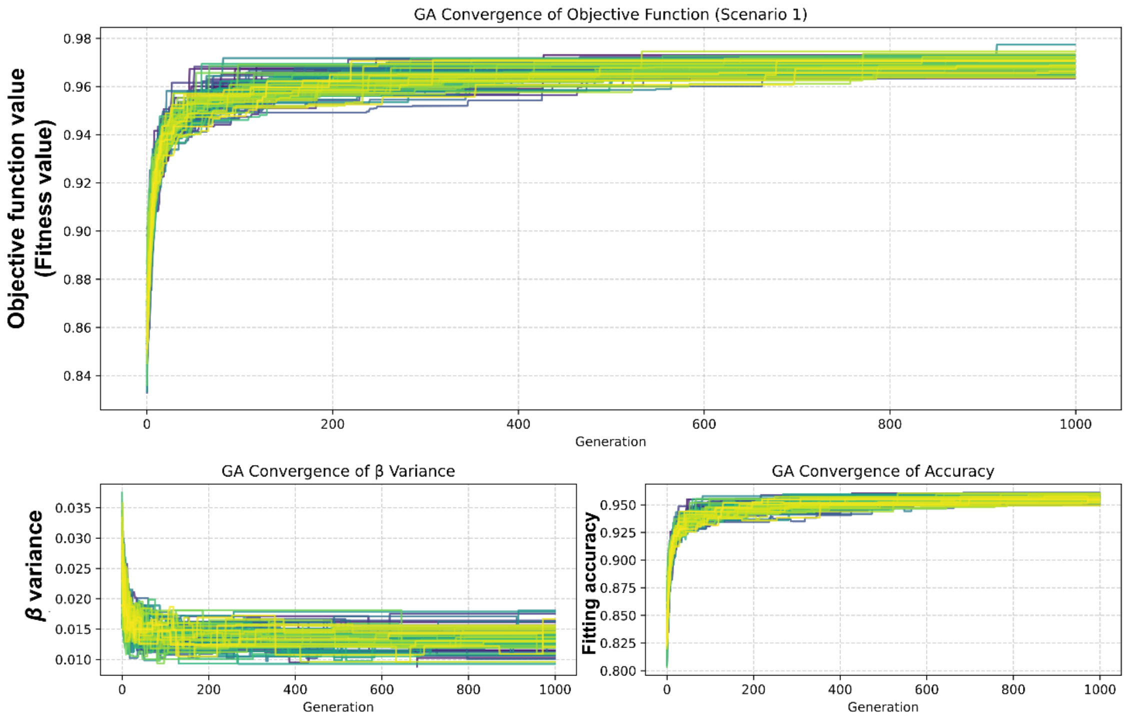


**Supplementary Figure 4. Convergence of the objective function with ELA/GAopt over 1000 generations for 100 runs with different starting population (Scenario 1, on HCP-YA data).** The objective function (top figure) is the sum of the β variance (bottom left) and the pMEM fitting accuracy (bottom right).

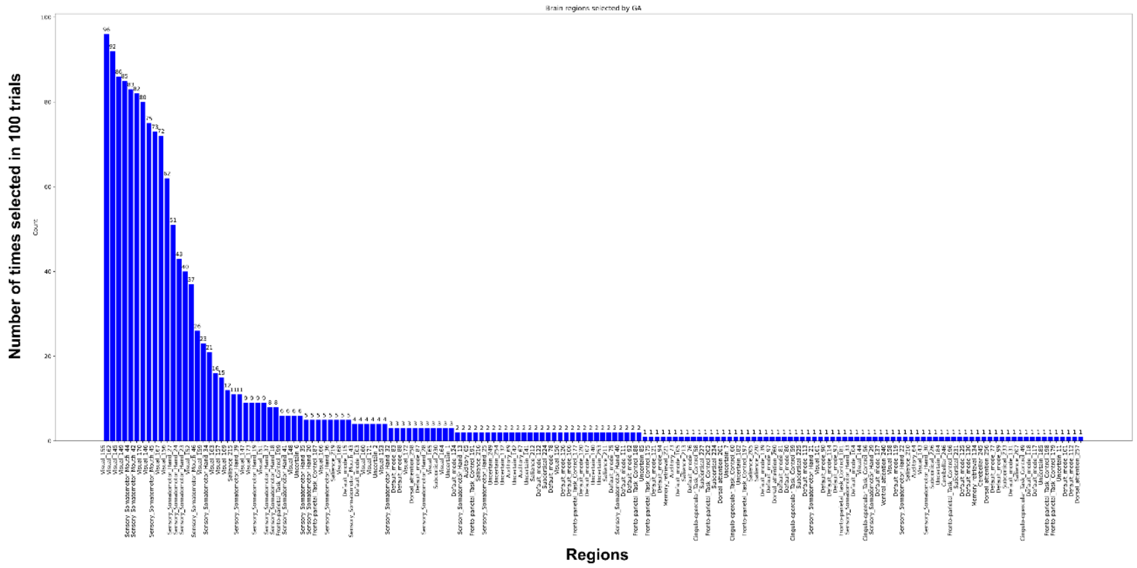


**Supplementary Figure 5. Selection frequency of each ROI across 100 runs of ELA/GAopt (Scenario 1, on HCP-YA data).**

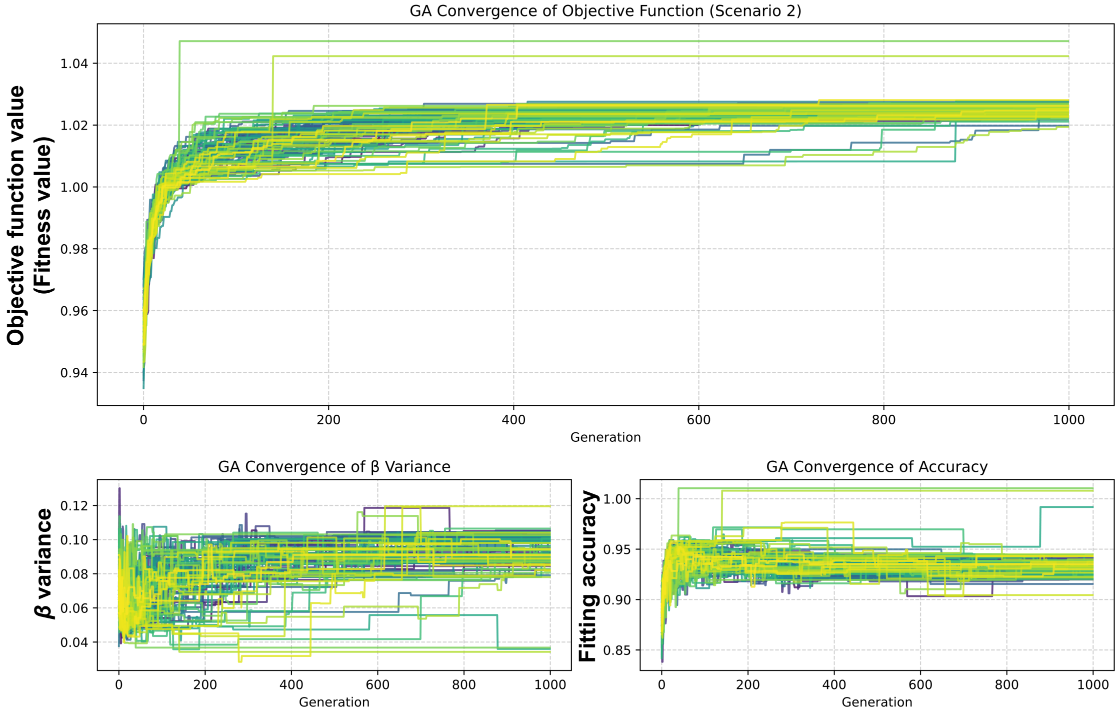


**Supplementary Figure 6. Convergence of the objective function with ELA/GAopt over 1000 generations for 100 runs with different starting population (Scenario 2).** The objective function (top figure) is the sum of the β variance (bottom left) and the pMEM fitting accuracy (bottom right).

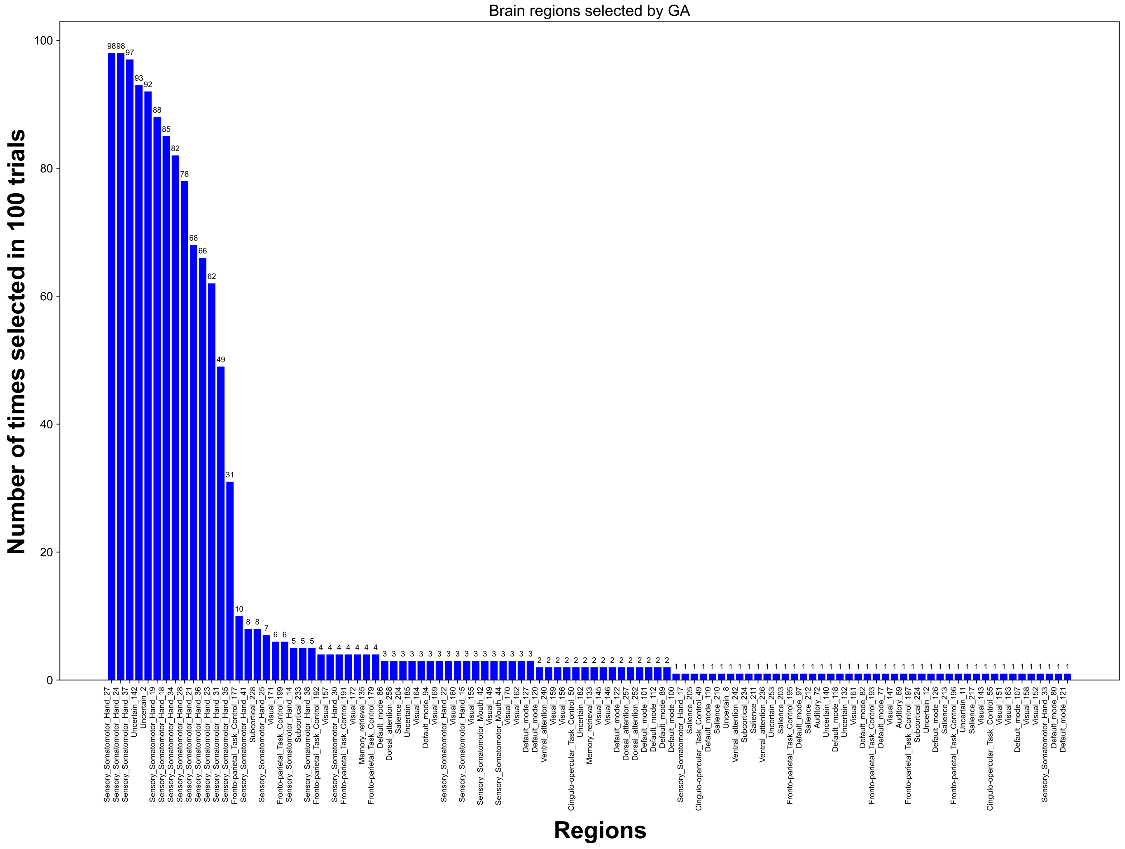


**Supplementary Figure 7. Selection frequency of each ROI across 100 runs of ELA/GAopt (Scenario 2).**

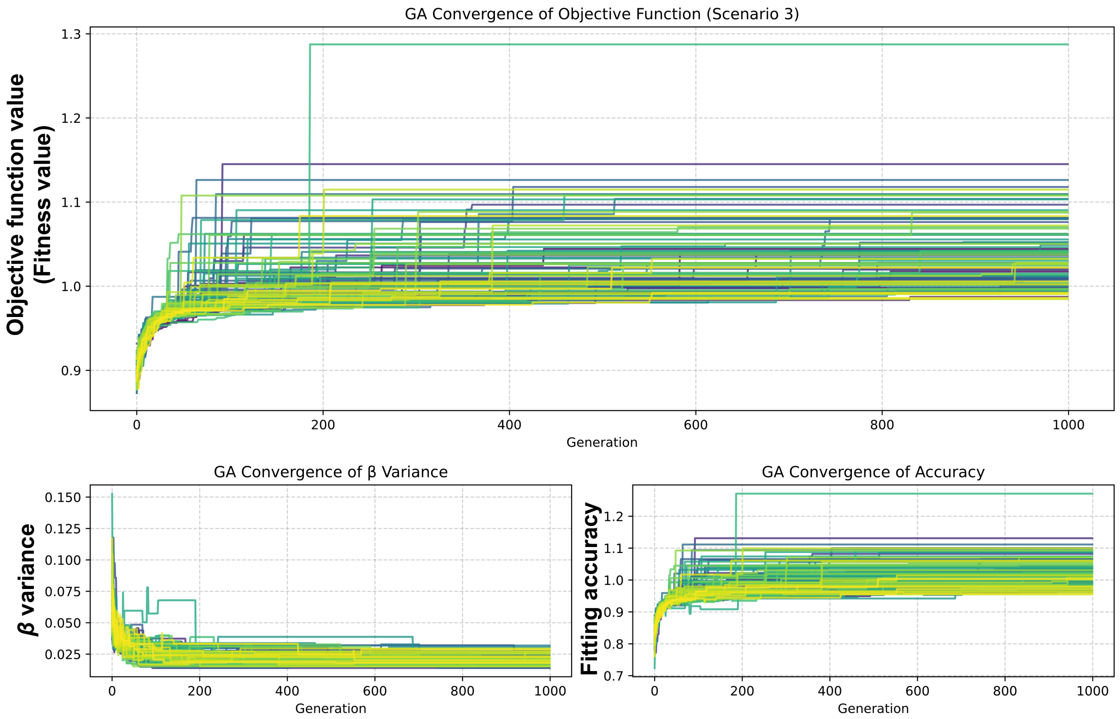


**Supplementary Figure 8. Convergence of the evaluation function across generations during 100 runs of ELA/GAopt (Scenario 3).** The objective function (top figure) is the sum of the β variance (bottom left) and the pMEM fitting accuracy (bottom right).

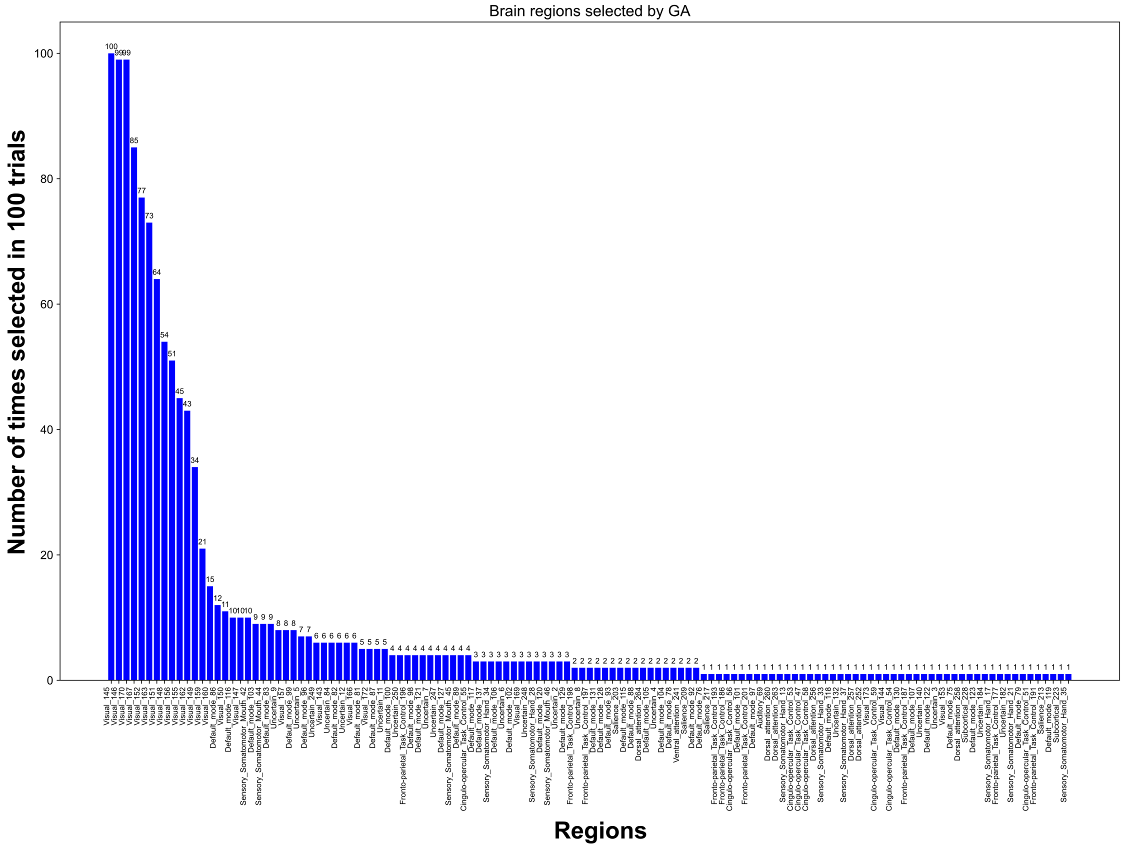


**Supplementary Figure 9. Selection frequency of each ROI across 100 runs of ELA/GAopt (Scenario 3).**

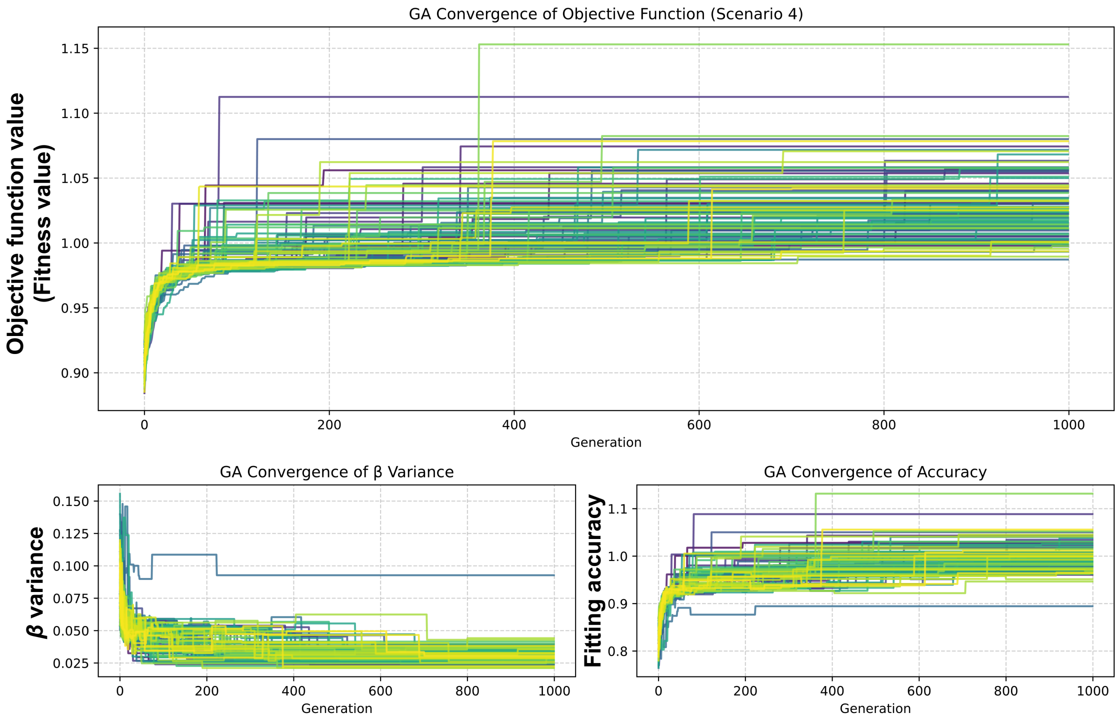


**Supplementary Figure 10. Convergence of the evaluation function across generations during 100 runs of ELA/GAopt (Scenario 4).** The objective function (top figure) is the sum of the β variance (bottom left) and the pMEM fitting accuracy (bottom right).

ELA with Genetic ROI Optimization

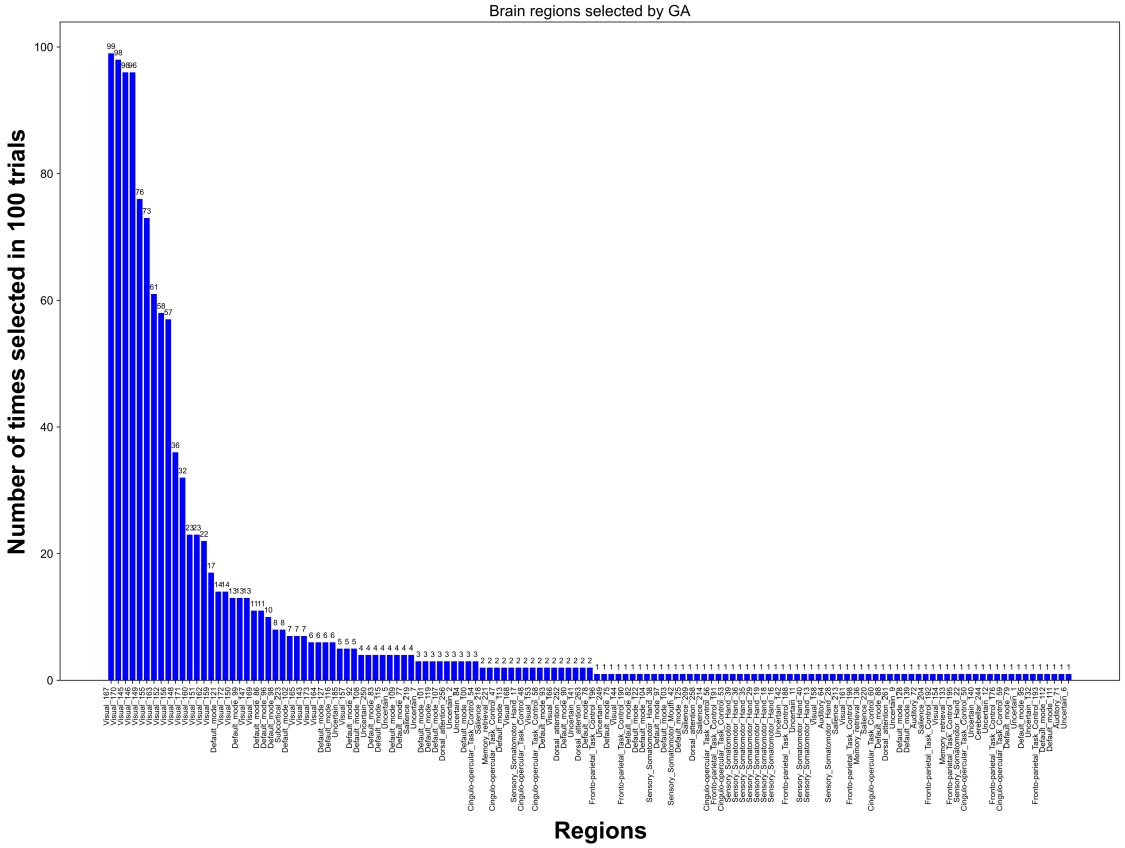


**Supplementary Figure 11. Selection frequency of each ROI across 100 runs of ELA/GAopt (Scenario 4).**
